# Combined forces of hydrostatic pressure and actin polymerization drive endothelial tip cell migration and sprouting angiogenesis

**DOI:** 10.1101/2024.05.20.594949

**Authors:** Igor Kondrychyn, Liqun He, Haymar Wint, Christer Betsholtz, Li-Kun Phng

## Abstract

Cell migration is a key process in the shaping and formation of tissues. During sprouting angiogenesis, endothelial tip cells invade avascular tissues by generating actomyosin-dependent forces that drive cell migration and vascular expansion. Surprisingly, ECs can still invade if actin polymerization is inhibited. In this study, we show that endothelial tip cells employ an alternative mechanism of cell migration that is dependent on Aquaporin (Aqp)-mediated water inflow and increase in hydrostatic pressure. In the zebrafish, ECs express *aqp1a.1* and *aqp8a.1* in newly formed vascular sprouts in a VEGFR2-dependent manner. Aqp1a.1 and Aqp8a.1 loss-of-function studies show an impairment in intersegmental vessels formation because of a decreased capacity of tip cells to increase their cytoplasmic volume and generate membrane protrusions, leading to delayed tip cell emergence from the dorsal aorta and slower migration. Further inhibition of actin polymerization resulted in a greater decrease in sprouting angiogenesis, indicating that ECs employ two mechanisms for robust cell migration *in vivo*. Our study thus highlights an important role of hydrostatic pressure in tissue morphogenesis.

## INTRODUCTION

The blood vascular system is a dynamic, multicellular tissue that adapts to the metabolic demands of development, growth and homeostasis by altering its pattern and morphology. Macroscopic transformation in the vascular network is achieved by microscopic changes in endothelial cell (EC) behaviours. For example, vascular expansion through sprouting angiogenesis requires the coordination of EC migration, proliferation, anastomosis, lumen formation and cell rearrangements, all of which require specific cell shape changes. At the initial phase of sprouting angiogenesis, endothelial tip cells generate numerous membrane protrusions such as filopodia and lamellipodia that drive migration and anastomosis (Figueiredo et al., 2021; Gerhardt et al., 2003; Phng et al., 2013; Sauteur et al., 2017). During tubulogenesis, apical membranes deform by generating inverse blebs to expand lumens so that blood can be transported efficiently through the blood vascular network (Gebala et al., 2016). However, following vessel perfusion, ECs become less dynamic and instead develop mechanoresistance to the deforming forces of blood pressure to maintain vessel morphology (Kondrychyn et al., 2020). EC shape is therefore dynamic and changes depending on the morphogenetic state of vascular development.

It is well established that localized remodelling of actin cytoskeleton and non-myosin II contractility are instrumental in driving cell shape changes during tissue morphogenesis (Clarke and Martin, 2021; Munjal and Lecuit, 2014; Murrell et al., 2015). More recently, there is an accumulating body of work implicating a role of water flow and the resulting increase in hydrostatic pressure as another mechanical mechanism of cell shape change (Choudhury et al., 2022; Chugh et al., 2022; Li et al., 2020). For example, in the osmotic engine model of cell migration, polarized distribution of ion channels and transporters generate an osmotic gradient that drives water inflow at the leading edge and outflow at the rear, leading to forward translocation of tumour cells in confined microenvironment (Stroka et al., 2014). Additionally, hydrostatic pressure from the extracellular environment can drive tissue morphogenesis, as demonstrated during mouse blastocyst formation (Chan et al., 2019; Dumortier et al., 2019), development of the otic vesicle in the zebrafish (Mosaliganti et al., 2019) and blood vessel lumen expansion (Gebala et al., 2016). Hydrostatic pressure can therefore generate sufficient forces to shape tissues and organs to their proper form and size during development.

We have previously discovered that EC migration persists after the inhibition of actin polymerization and in the absence of filopodia to generate new blood vessels in the zebrafish (Phng et al., 2013). This observation suggests the existence of an alternative mechanism of migration independent of actin polymerization. In this study, we sought to determine whether hydrostatic pressure regulates EC migration during sprouting angiogenesis in the zebrafish by investigating the function of Aquaporins (Aqp), which are transmembrane water channels that increase water permeability of cell membranes to promote transcellular water flow (Day et al., 2014; Farinas et al., 1997; Kozono et al., 2002; Preston et al., 1992). We discovered that ECs of newly formed intersegmental vessels (ISV) express *aqp1a.1* and *aqp8a.1* mRNA, and observed the enrichment of Aqp1a.1 and Aqp8a.1 proteins in the leading edge of migrating tip cells. Detailed single cell analyses showed that endothelial tip cells lacking *aqp1a.1* and *aqp8a.1* expression have reduced cell volume, membrane protrusions and migration capacity. As a result, there is defective sprouting angiogenesis and ISV formation in *aqp1a.1;aqp8a.1* double mutant zebrafish. Notably, when actin polymerization is inhibited in *aqp1a.1*;*aqp8a.1* double mutant zebrafish, there is a greater impairment in EC migration and sprouting angiogenesis, demonstrating the additive function of actin polymerization and hydrostatic pressure (that is elevated by water inflow) in generating membrane protrusions to drive EC migration *in vivo*.

## RESULTS

### *aqp1a.1* and *aqp8a.1* are differentially expressed by tip and stalk cells during sprouting angiogenesis

Of the 18 *aquaporin* genes that are expressed in zebrafish (Tingaud-Sequeira et al., 2010), *aqp1a.1* and *aqp8a.1* mRNA have been detected in blood endothelial cells (ECs) (Koun et al., 2016; Rehn et al., 2011). We confirmed the endothelial expression of *aqp1a.1* and *aqp8a.1* at 30 hours post fertilization (hpf) and 2-, 3- and 4-days post fertilization (dpf) by whole-mount *in situ* hybridization (WISH, Fig. S1). The two endothelial *aqp* genes, however, differ in spatial expression. While *aqp1a.1* is widely expressed in blood vessels of the head and trunk (dorsal aorta (DA), caudal artery (CA), ISVs, dorsal longitudinal anastomotic vessels (DLAV) and caudal vein plexus) at 30 hpf, *aqp8a.1* expression is absent in cerebral blood vessels and is restricted to the DA, CA and the ventral regions of ISVs in the trunk. After 1 dpf, *aqp8a.1* expression expands to the entire ISV and DLAV. Additionally, the expression of *aqp1a.1* and *aqp8a.1* gradually decreases in the DA and CA so that both are absent by 4 dpf. Single cell RNA sequencing (scRNAseq) of ECs isolated at 24 hpf, 34 hpf and 3 dpf further confirmed the endothelial expression of *aqp1a.1* and *aqp8a.1* mRNA (Fig. S2 A - F), revealed differential expression of *aqp1a.1* and *aqp8a.1* mRNA in distinct endothelial clusters and highlighted higher *aqp1a.1* transcript expression in all endothelial subtypes compared to *aqp8a.1* (Fig. S2 E - H).

We next examined the expression pattern of *aqp1a.1* and *aqp8a.1* at the beginning of sprouting angiogenesis at higher resolution using RNAscope and confocal microscopy. At 20 hpf, *aqp1a.1* mRNA is highly expressed in the DA with little expression in the PCV (Fig. 1A, C and Fig. S3). Closer inspection uncovered heterogeneous expression in the DA, with higher expression in ECs that would be specified as tip cells (Fig. 1A, C and Fig. S3B). Indeed, tip cells of newly formed vascular sprouts express higher levels of *aqp1a.1* compared to adjacent ECs in the DA at 20 hpf (Fig. 1A – C, Fig. S3B) and the majority of ISVs (61%) display higher *aqp1a.1* expression in tip cells than in trailing stalk cells at 22 hpf (Fig. 1G). *Aqp8a.1* expression is at first largely absent in the DA at 20 hpf (Fig. 1B, D and Fig. S3C) but becomes detectable at 22hpf (Fig. 1E and F). Interestingly, unlike *aqp1a.1*, *aqp8a.1* expression is undetectable in tip cells of emerging ISVs at 20 hpf. At 22 hpf, it is expressed in 22 out of 23 tip cells analyzed with 78% of ISVs exhibiting higher *aqp8a.1* mRNA expression in stalk cells than tip cells (Fig. 1G). These observations highlight differential expression pattern of *aqp1a.1* and *aqp8a.1* in newly formed vascular sprouts, with *aqp1a.1* expression enriched in tip cells and *aqp8a.1* expression in stalk cells.

**Figure 1.**
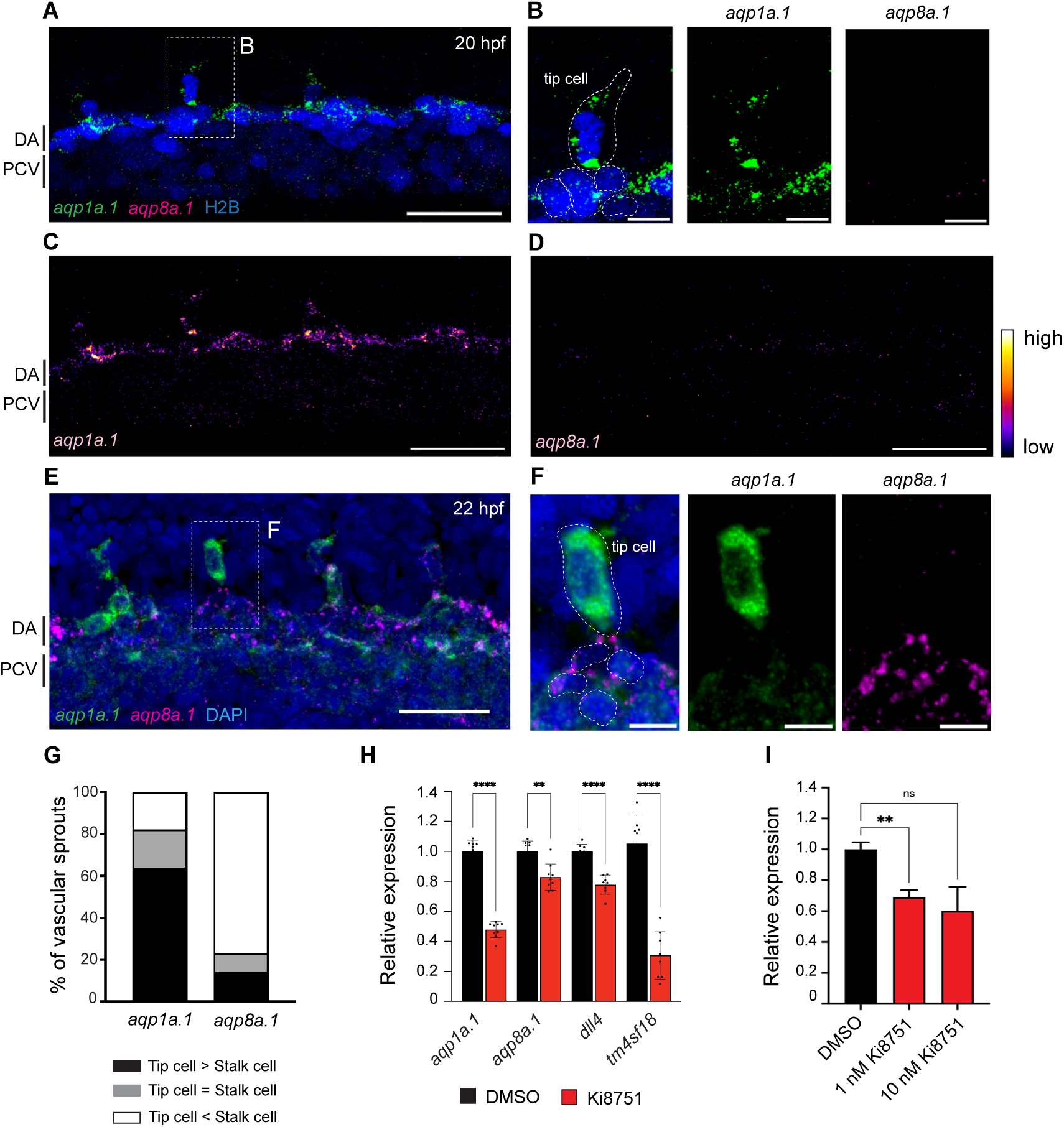
Differential expression of *aqp1a.1* and *aqp8a.1* mRNAs in tip and stalk cells. (**A-F**) Detection of *aqp1a.1* and *aqp8a.1* mRNA by RNAscope *in situ* hybridization in 20 (A - D) and 22 (E and F) hpf embryos. Representative maximum intensity projection of confocal z-stacks of tip and stalk cells sprouting from DA are shown (n = 9 embryos, 2 independent experiments). B and F show magnified region of a tip cell from a 20 and 22 hpf zebrafish, respectively (tip cell and EC nuclei in the DA are outlined). Scale bar, 10 µm (B and F) and 40 µm (A, C - E). (**G**) Percentage of vascular sprouts with differential expression of *aqp1a.1* and *aqp8a.1* mRNA in tip and stalk cells in 22 hpf embryos (n=23 sprouts from 8 embryos, 2 independent experiments). (**H**) qPCR analysis of *aqp1a.1* and *aqp8a.1* expression in zebrafish embryos treated with 0.01% DMSO or 1 µM ki8751 at 20 hpf for 6 hours (n = 3 independent experiments). (**I**) qPCR analysis of *AQP1* gene expression in HAECs treated with 0.01% DMSO, 1 or 10 nM ki8751 for 6 hours (n = 2 independent experiments). In panels H and J, gene expressions are shown relative to *gapdh* expression and data are presented as mean ± SD; statistical significance was determined by Brown-Forsythe and Welch ANOVA tests with Dunnett’s multiple comparisons test; ns *p*>0.05, *\*\*p*<0.01, *\*\*\*p*<0.001, and *\*\*\*\*p*<0.0001. DA, dorsal aorta; PCV, posterior cardinal vein. See also Fig. S1-S3.

### VEGF-VEGFR2 signalling induces the expression of *aqp1a.1* and *aqp8a.1* mRNA

We next explored whether *aqp1a.1* and *aqp8a.1* expression is regulated by VEGFR2 signalling, which is highly activated in tip cells during sprouting angiogenesis (Gerhardt et al., 2003; Siekmann et al., 2013). To inhibit VEGFR2 activity, embryos were treated with 1 µM ki8751 from 20 hpf. Relative quantitative polymerase chain reaction (qPCR) experiments showed that *aqp1a.1* and *aqp8a.1* mRNA expression was suppressed by 52% (*p* < 0.0001) and 17% (*p* = 0.0028), respectively, after 6 hours of ki8751 treatment when compared to DMSO-treated embryos (Fig. 1H). These results demonstrate that VEGFR2 activity upregulates the expression of *aqp1a.1* and *aqp8a.1*. To examine whether VEGFR2 activity also regulates Aquaporin expression in human ECs, we treated cultured human aortic endothelial cells (HAECs) with ki8751 for 6 hours and analyzed *AQP1* expression by qPCR. *AQP8* expression was not examined since it is not expressed in HAECs (data not shown). Inhibition of VEGFR2 activity with 1 nM and 10 nM ki8751 decreased *AQP1* mRNA expression by 31% (*p* = 0.0023, Fig. 1I) and 40% (*p* = 0.1032, Fig. 1I), respectively, demonstrating cross-species conservation of VEGFR2 activity in inducing *Aqp1* mRNA expression.

### Aqp1a.1 and Aqp8a.1 proteins are enriched at the leading edge of migrating tip cells

To gain a better understanding of where within the cell Aquaporin protein functions, we generated stable *Tg(fli1ep:aqp1a.1-mEmerald)^rk30^* and *Tg(fli1ep:aqp8a.1-mEmerald)^rk31^*zebrafish lines in which ECs express Aqp1a.1 or Aqp8a.1 tagged with mEmerald, respectively. At 25 hpf, we observed an accumulation of Aqp1a.1 (Fig. 2A) and Aqp8a.1 (Fig. 2B) in mCherryCAAX-positive membrane clusters in the cytoplasm and filopodia of tip cells. Notably, many of the Aqp1a.1- or Aqp8a.1-positive membrane clusters are located at the migrating front of the tip cell. Time-lapse imaging capturing the process of ISV formation between 24 to 30 hpf further showed sustained enrichment of Aqp1a.1 and Aqp8a.1 in clusters at the front of the tip cell as well as in filopodia (Fig. 2C and D, Movie S1 and S2), suggesting that Aqp1a.1 and Aqp8a.1 may promote localized water flux at the migrating edge of tip cells.

**Figure 2.**
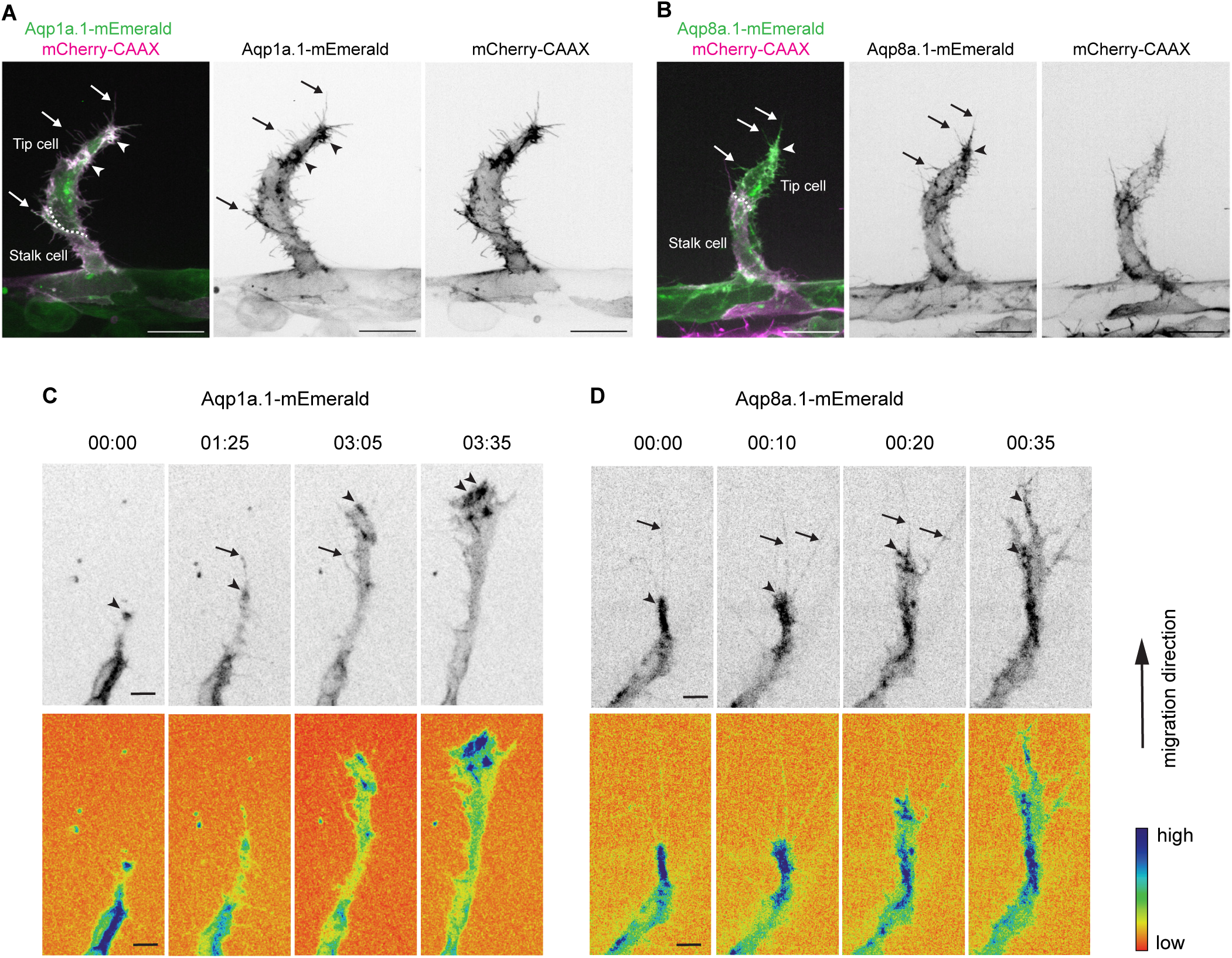
Aquaporin proteins are enriched at the leading edge of migrating tip cells. (**A-B**) Representative maximum intensity projection confocal z-stacks of endothelial tip cells of 25 hpf *Tg(fli1ep:aqp1a.1-mEmerald)^rk30^;(kdrl:ras-mCherry)^s916^*(A) and *Tg(fli1ep:aqp8a.1-mEmerald)^rk31^;(kdrl:ras-mCherry)^s916^*(B) embryos. White serrated line outlines tip-stalk cell border. Scale bar, 20 µm. (**C-D**) Still images from time-lapse movies of migrating tip cells from *Tg(fli1ep:aqp1a.1-mEmerald)^rk30^* (C) and *Tg(fli1ep:aqp8a.1-mEmerald)^rk31^* (D) embryos. Movies were taken from 24 to 30 hpf. Arrowheads, Aquaporin protein localization at the leading edge of migrating tip cells. Arrows, Aquaporin localization in filopodia. Time, h:min. Scale bar, 5 µm. See also Movies S1 and S2.

### Loss of Aqp1a.1 and Aqp8a.1 function leads to defective trunk blood vessel formation

We subsequently proceeded to generate zebrafish *aqp1a.1* and *aqp8a.1* mutants to investigate the function of Aquaporin-mediated water flow in blood vessel development. Using CRISPR/Cas9 genome editing, we targeted the first exons of *aqp1a.1* and *aqp8a.1* genes. The CRISPR-generating allele *aqp1a.1^rk28^* carries a 14-bp deletion and an 8-bp insertion in the 5’ end of the gene leading to a premature termination codon at amino acid 76 after 1 missense amino acid (Fig. S4A-B). The *aqp8a.1^rk29^* allele carries a 4-bp deletion in the 5’ end of the gene that leads to a frameshift after Thr15 and premature termination codon at amino acid 73 after 58 missense amino acids (Fig. S5A-B). Using whole-mount *in situ* hybridization and qPCR analysis, we demonstrated that *aqp1a.1* and *aqp8a.1* mRNA expression is significantly decreased in *aqp1a.1^rk28/rk28^* (Fig. S4C-D) and *aqp8a.1^rk29/rk29^* (Fig. S5C-D) zebrafish embryos, respectively, suggesting a rapid degradation of mutant mRNAs and as a result, the loss of Aquaporin protein expression.

Although *aqp1a.1^rk28/rk28^;aqp8a.1^rk29/rk29^*embryos look morphologically normal when compared to wildtype or *aqp1a.1^+/rk28^;aqp8a.1^+/rk29^*embryos (Fig. S8A-D), we observed defects in blood vessel formation and patterning at 1 to 3 dpf. Compared to 28 hpf control embryos whose ISVs have reached the dorsal roof of the neural tube and started to form the DLAV, ISVs of *aqp1a.1^rk28/rk28^;aqp8a.1^rk29/rk29^*embryos are much less developed (Fig. 3A and D) with significantly reduced length (Fig. 3E and Fig. S6H and K). Single deletion of *aqp1a.1* (Fig. 3B and Fig. S6F and I) or *aqp8a.1* (Fig. 3C and Fig. S6G and J) also led to defective ISV formation, although the defects are less severe compared to the loss of both *aqp1a.1* and *aqp8a.1* expression. Comparison of ISV length in *aqp1a.1^rk28/rk28^*, *aqp8a.1^rk29/rk29^* and *aqp1a.1^rk28/rk28^;aqp8a.1^rk29/rk29^*embryos shows that the loss of *aqp1a.1* leads to a greater decrease in ISV length than the loss of *aqp8a.1* (Fig. 3E, Fig. S6I and J), and that the loss of both *aqp1a.1* and *aqp8a.1* results in the greatest reduction. These observations indicate an additive effect of Aqp1a.1 and Aqp8a.1 function in blood vessel development and implicate Aqp1a.1 as the more dominant Aquaporin protein in EC function, in line with the higher expression of *aqp1a.1* mRNA in tip cells.

**Figure 3.**
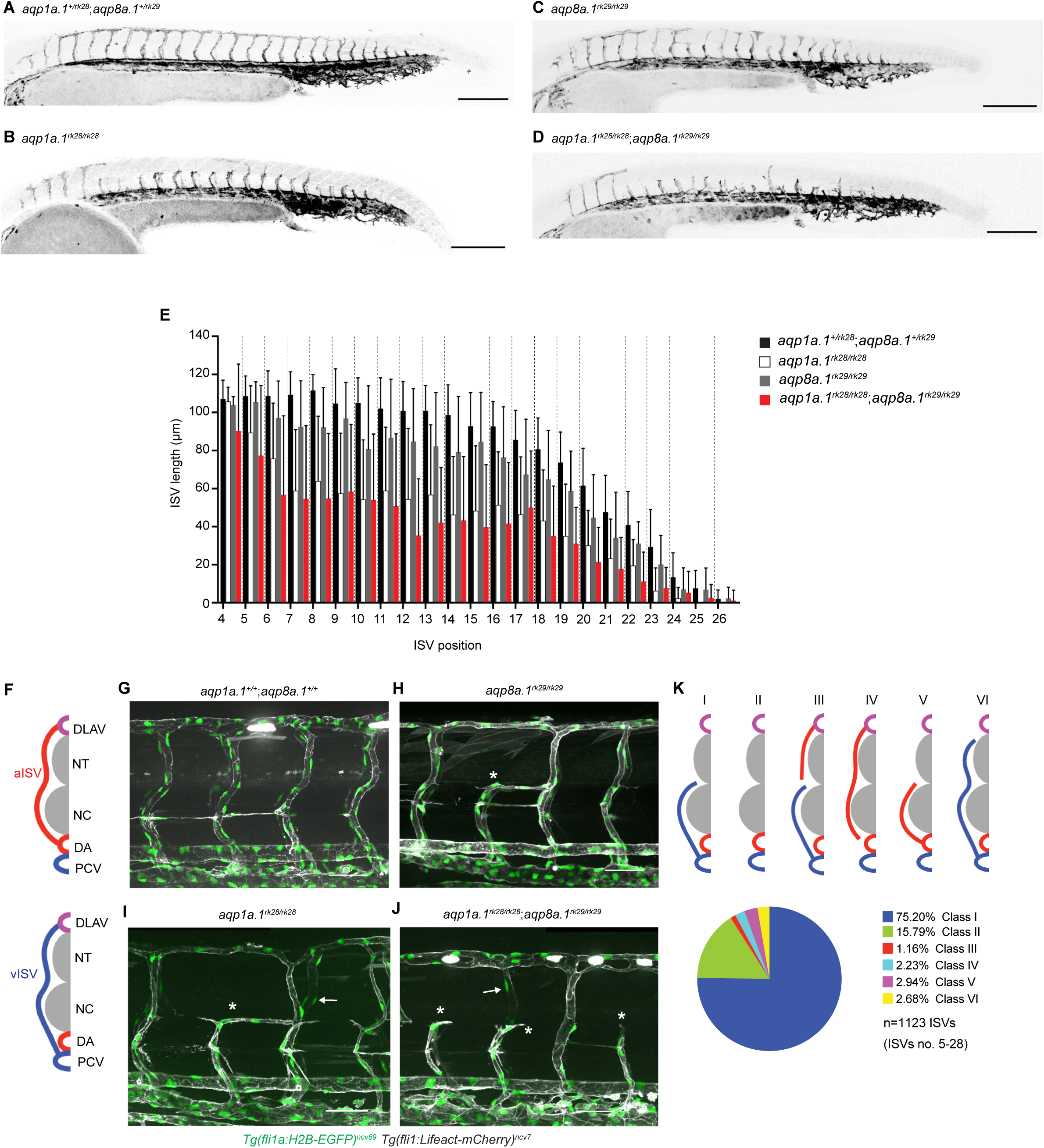
Loss of Aquaporin function leads to defective trunk vessel formation. (**A-D**) Representative maximum intensity projection confocal z-stacks of 28 hpf *aqp1a.1^+/rk28^;aqp8a.1^+/rk29^* (A), *aqp1a.1^rk28/rk28^*(B), *aqp8a.1^rk29/rk29^* (C) and *aqp1a.1^rk28/rk28^;aqp8a.1^rk29/rk29^*(D) embryos. (**E**) Quantification of ISV length at different positions along the trunk of 28 hpf embryos (*aqp1a.1^+/rk28^;aqp8a.1^+/rk29^*: n = 21 embryos; *aqp1a.1^rk28/rk28^*: n = 20 embryos; *aqp8a.1^rk29/rk29^*: n = 20 embryos; *aqp1a.1^rk28/rk28^;aqp8a.1^rk29/rk29^*: n = 19 embryos from 2 independent experiments). (**F**) Illustration of aISV and vISV connections in the trunk of wild type embryo at 3 dpf. (**G-J**) Representative maximum intensity projection confocal z-stacks of 3 dpf wild type (G), *aqp8a.1^rk29/rk29^* (H), *aqp1a.1^rk28/rk28^*(I), and *aqp1a.1^rk28/rk28^;aqp8a.1^rk29/rk29^* (J) embryos. Asterisk marks incomplete ISVs, arrows point EC nuclei on the contralateral side. (**K**) Pie chart showing proportion of different classes of truncated ISV phenotypes (I-VI) found in *aqp1a.1^rk28/rk28^;aqp8a.1^rk29/rk29^* mutant embryos at 3 dpf. DA, dorsal aorta; DLAV, dorsal longitudinal anastomotic vessel; ISV, intersegmental vessel; aISV, arterial ISV; vISV, venous ISV; NC, notochord; NT, neural tube; PCV, posterior cardinal vein. Scale bar, 50 µm (G-J) and 200 µm (A-D). See also Fig. S6-S8.

We next examined the embryos at 3 dpf, a stage when arterial ISVs (aISVs) and venous (vISVs) are fully established and lumenized, connecting the DLAV to either the DA or PCV (Fig. 3F and G), respectively. Such fully established ISVs are significantly reduced in embryos when *aqp1a.1*, *aqp8a.1* or both *aqp1a.1* and *aqp8a.1* are deleted. Instead, these embryos display an increased number of aISVs and vISVs that fail to establish a connection to the DLAV, DA or PCV (Fig. 3H – J). The number of incompletely formed or truncated ISVs per embryo in *aquaporin* mutants was variable, between 1 to 6, 1 to 16 and 1 to 22 truncated ISVs in *aqp8a.1^rk29/rk29^*, *aqp1a.1^rk28/rk28^* and *aqp1a.1^rk28/rk28^;aqp8a.1^rk29/rk29^*embryos, respectively (Fig. S6A-D). We also found that truncated ISVs appear with higher frequency in the zebrafish trunk (ISV number 5 to 15) rather than in the tail (ISV number 16 to 27, Fig. S6E). Detailed analysis revealed that most of the truncated ISVs are vISVs that are connected to the PCV but not the DLAV (Class I in Fig. 3K), followed by the absence of aISV or vISV (Class II in Fig. 3K, Fig. S7A and B). A small fraction of the truncated vessels are aISVs that are not connected to the DA (Class IV in Fig. 3K, Fig. S7D) or DLAV (Class V in Fig. 3K, Fig. S7E), or vISVs that are not connected to the DLAV (Class VI in Fig. 3K, Fig. S7F). At some regions, there is a truncated aISV that is not connected to the DA and an incomplete vISV that is not connected to the DLAV (Class III in Fig. 3K, Fig. S7C).

As *aqp1a.1* is expressed in the brain vasculature during development (Fig. S1), we examined whether cerebral vasculature is also affected in *aqp1a.1^rk28/rk28^* embryos. At 4 dpf, we found that the length of the dorsal longitudinal vein (DLV), a main blood vessel that supplies blood to the choroid plexus (Bill and Korzh, 2014) was reduced in *aqp1a.1^rk28/rk28^* embryos (*p* = 0.0098, Fig. S9A, B and E). Also, in 2 out of 21 *aqp1a.1^rk28/rk28^* embryos, the DLV was undeveloped (Fig. S9C, D and F), a defect that is also observed in 50% of *vegfab^-/-^* and *vegfc^-/-^* mutant zebrafish (Parab et al., 2021).

In summary, our analyses show that in the absence of Aqp1a.1 and/or Aqp8a.1 function, there is an initial delay in ISV formation during primary sprouting angiogenesis. By 3 dpf, some ISVs are completely formed (Fig. S8K and M), connecting the DA or PCV to the DLAV. However, the number of completely formed ISVs is significantly decreased, with the loss of Aqp1a.1 having a more severe impact compared to the loss of Aqp8a.1, and the combined loss of Aqp1a.1 or Aqp8a.1 having a greater deleterious effect.

### Aquaporins promote endothelial tip cell protrusion and migration

To gain a better understanding of why ISVs do not develop normally, we assessed EC number. By counting the number of nuclei in aISVs and vISVs, we found a significant reduction in EC number per vessel in 2 and 3 dpf *aqp1a.1^rk28/rk28^;aqp8a.1^rk29/rk29^* embryos when compared to wildtype embryos (Fig. S10A and B), suggesting that the defective formation of ISVs may result from insufficient EC number. Quantification of endothelial mitotic events between 21 and 30 hpf shows that there is a similar number of tip and stalk cell division in *aqp1a.1^+/rk28^;aqp8a.1^+/rk29^*embryos and *aqp1a.1^rk28/rk28^;aqp8a.1^rk29/rk29^* embryos (Fig. S10C), indicating that decreased EC division is not the cause of reduced EC number in ISVs. We next explored whether EC migration from the DA is impaired by examining tip cell behaviour from 21 – 22hpf, when sprouting angiogenesis begins. In *aqp1a.1^+/rk28^;aqp8a.1^+/rk29^* embryos, tip cells emerge from the DA between 20 and 24 hpf (Fig. 4A) and migrate dorsally between the somite boundaries over the notochord and neural tube to form primary ISVs (Fig. 4B, Movie S3). However, in *aqp1a.1^rk28/rk28^;aqp8a.1^rk29/rk29^*embryos, tip cell nuclei emerge between 22 and 28 hpf (Fig. 4A). Furthermore, in some *aqp1a.1^rk28/rk28^;aqp8a.1^rk29/rk29^* embryos, tip cell membrane protrusions do not extend dorsally in the direction of migration but retract back to the DA so that a primary ISV does not form at this location (Fig. 4C, Movie S4), potentially giving rise to the Class II phenotype at 3 dpf (Fig. 3K and Fig. S7B). These observations indicate that tip cell protrusion from the DA is significantly delayed in the absence of Aqp1a.1 and Aqp8a.1 function.

**Figure 4.**
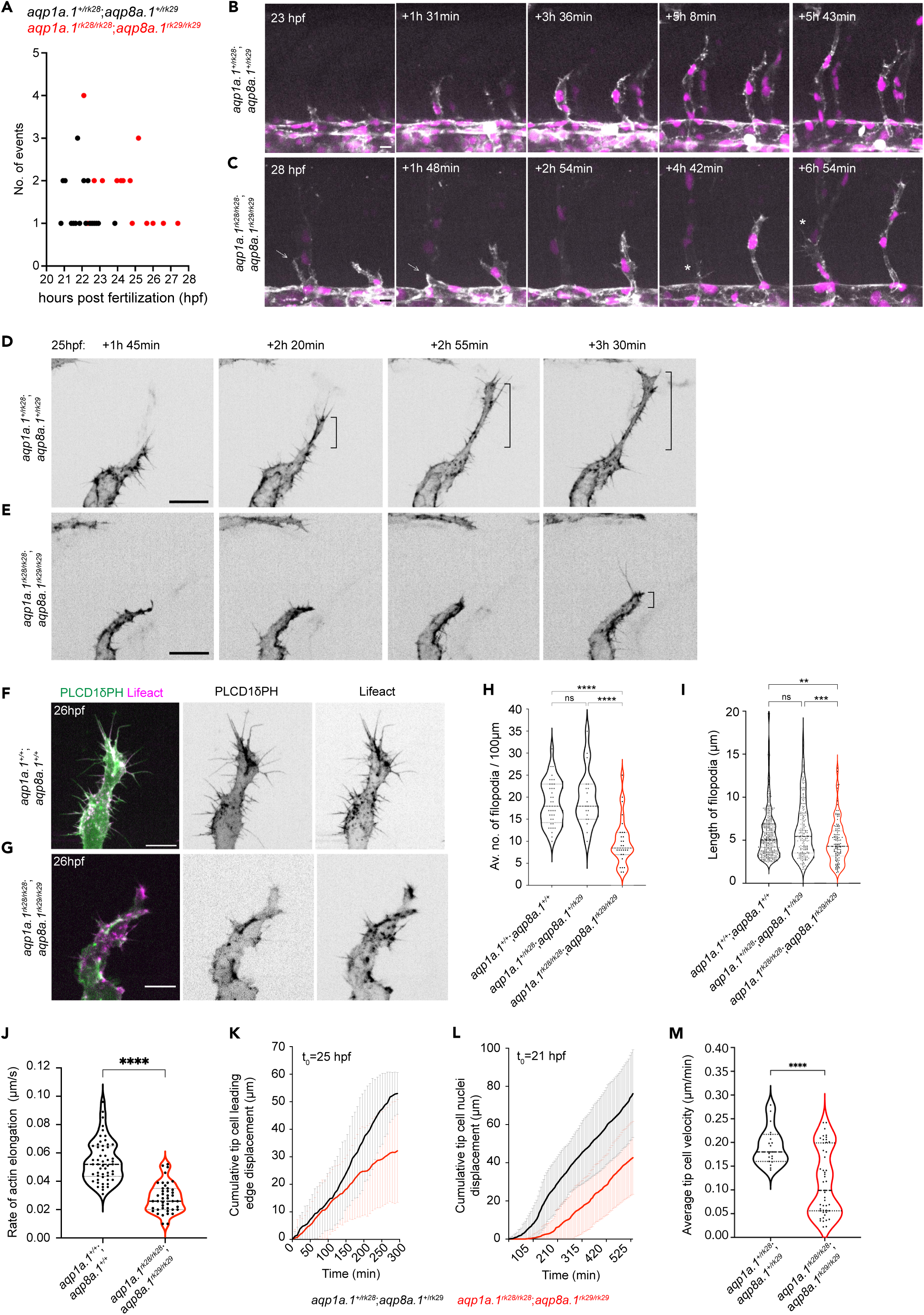
Aquaporins promote endothelial tip cell protrusion and migration. (**A**) Timing of tip cell emergence from the DA in *aqp1a.1^+/rk28^;aqp8a.1^+/rk29^* (n = 23 cells) and *aqp1a.1^rk28/rk28^;aqp8a.1^rk29/rk29^* (n = 27 cells). (**B-C**) Still images from time-lapse imaging of migrating tip cells from *aqp1a.1^+/rk28^;aqp8a.1^+/rk29^* (B) and *aqp1a.1^rk28/rk28^;aqp8a.1^rk29/rk29^* (C) embryos from 20 to 30 hpf. White arrows, retracting tip cell. *, secondary sprouting from PCV. Scale bar, 10 µm. (**D-E**) Stills images from representative time-lapse movies of migrating tip cells in *aqp1a.1^+/rk28^;aqp8a.1^+/rk29^* (D, n = 2) and *aqp1a.1^rk28/rk28^;aqp8a.1^rk29/rk29^* (E, n = 5) embryos. Movies were taken from 25 to 30 hpf (n = 2 independent experiments). Bracket, formation of stable protrusions. Scale bar, 20 µm. (**F-G**) Representative maximum intensity projection confocal z-stacks of tip cells from wild type (F) and *aqp1a.1^rk28/rk28^;aqp8a.1^rk29/rk29^* (G) embryos at 26 hpf. Scale bar, 10 µm. (**H**) Quantification of filopodia number in tip cells of wild type (n = 36 cells from 10 embryos, 2 independent experiments), *aqp1a.1^+/rk28^;aqp8a.1^+/rk29^* (n = 19 cells from 6 embryos, 2 independent experiments) and *aqp1a.1^rk28/rk28^;aqp8a.1^rk29/rk29^* (n = 28 cells from 9 embryos, 2 independent experiments) embryos. (**I**) Quantification of filopodia length in tip cells of wild type (n = 24 cells from 7 embryos, 2 independent experiments), *aqp1a.1^+/rk28^;aqp8a.1^+/rk29^* (n = 16 cells from 7 embryos, 2 independent experiments) and *aqp1a.1^rk28/rk28^;aqp8a.1^rk29/rk29^* (n = 11 cells from 6 embryos, 2 independent experiments) embryos. (**J**) Quantification of growth rate of actin bundles in tip cell filopodia in 25 hpf wild type (n = 12 cells from 7 embryos, 2 independent experiments) and *aqp1a.1^rk28^;aqp8a.1^rk29^* (n = 12 cells from 6 embryos, 2 independent experiments) embryos. (**K**) Quantification of tip cell leading edge displacement of *aqp1a.1^+/rk28^;aqp8a.1^+/rk29^* (n = 19 cells from 5 embryos, 3 independent experiments) and *aqp1a.1^rk28/rk28^;aqp8a.1^rk29/rk29^* (n = 47 cells from 13 embryos, 6 independent experiments) embryos at 25 - 30 hpf. (**L**) Quantification of tip cell nuclei displacement of *aqp1a.1^+/rk28^;aqp8a.1^+/rk29^* (n = 20 cells from 6 embryos, 4 independent experiments) and *aqp1a.1^rk28/rk28^;aqp8a.1^rk29/rk29^*(n = 20 cells from 4 embryos, 2 independent experiments) embryos at 21 - 30 hpf. (**M**) Quantification of tip cell migration velocity in *aqp1a.1^+/rk28^;aqp8a.1^+/rk29^*(n = 19 cells from 5 embryos, 3 independent experiments) and *aqp1a.1^rk28/rk28^;aqp8a.1^rk29/rk29^*(n = 47 cells from 13 embryos, 6 independent experiments) embryos at 25 - 30 hpf. Statistical significance was determined by Brown-Forsythe and Welch ANOVA tests with Dunnett’s (H) or Sidak’s (I) multiple comparisons test, and with unpaired *t*-test (J, M). ns, *p*>0.05, **, *p* < 0.01, *\*\*\**, *p* < 0.001, and *\*\*\*\**, *p* < 0.0001. See also Fig. S10, Movies S3-6.

We have previously demonstrated that filopodia facilitate EC migration by serving as a template for lamellipodia formation (Phng et al., 2013). Lamellipodia frequently emanate laterally from stable filopodia at the leading edge of tip cells, leading to rapid expansion of the filopodium, increase in volume and the formation of a stable protrusion in the direction of migration (Phng et al., 2013). This was similarly observed in *aqp1a.1^+/rk28^;aqp8a.1^+/rk29^* embryos (Fig. 4D and Movie S5) but not in *aqp1a.1^rk28/rk28^;aqp8a.1^rk29/rk29^* embryos (Fig. 4E and Movie S6). Comparison of filopodia dynamics in wildtype (Fig. 4F), *aqp1a.1^+/rk28^;aqp8a.1^+/rk29^*and *aqp1a.1^rk28/rk28^;aqp8a.1^rk29/rk29^*(Fig. 4G) embryos revealed a significant reduction in filopodia number (wildtype, 18.73 ± 5.1/100µm; *aqp1a.1^+/rk28^;aqp8a.1^+/rk29^*, 20.4 ± 6.5/100µm; *aqp1a.1^rk28/rk28^;aqp8a.1^rk29/rk29^*, 9.8 ± 5.5/100µm; Fig. 4H) and length (wildtype, 5.5 ± 2.8 µm; *aqp1a.1^+/rk28^;aqp8a.1^+/rk29^*, 6.0 ± 3.4 µm; *aqp1a.1^rk28/rk28^;aqp8a.1^rk29/rk29^*, 4.5 ± 2.2 µm; Fig. 4I) in ECs lacking both Aqp1a.1 and Aqp8a.1 function that can be accounted for by a decrease in actin polymerization (Fig. 4J). By tracking Lifeact-mCherry signal in filopodia, we determined that the average elongation rate of actin as 0.055 ± 0.013 µm per second in wildtype embryos and that this is significantly decreased by 47% to 0.029 ± 0.011 µm per second (*p* < 0.0001) in *aqp1a.1^rk28/rk28^;aqp8a.1^rk29/rk29^*embryos. Analysis of time-lapse images of tip cell behaviour further showed slower tip cell membrane expansion at the leading edge (Fig. 4K), decreased nuclei displacement (Fig. 4L) and a significant reduction in tip cell migration velocity (Fig. 4M) in *aqp1a.1^rk28/rk28^;aqp8a.1^rk29/rk29^* embryos. To determine whether the slower EC migration is caused by a loss of front-rear polarity, we examined Golgi position relative to the nucleus in wildtype, *aqp1a.1^rk28/rk28^* and *aqp8a.1^rk29/rk29^*embryos at 24 - 26 hpf (Fig. S10D). This analysis shows that tip cell polarity is unperturbed in the absence of Aqp1a.1 or Aqp8a.1 function.

In conclusion, our findings demonstrate that Aqp1a.1 and Aqp8a.1 regulate sprouting angiogenesis by promoting endothelial tip cell emergence from the DA, the formation and expansion of membrane protrusions and EC migration.

### Tip cell volume regulation depends on Aquaporin-mediated water influx

Aquaporin channels permit bidirectional flow of water across the plasma membrane, with the direction of flow determined by the osmotic gradient between the extracellular environment and cell cytoplasm (Agre et al., 2002). We next sought to determine whether water flows in or out of tip cells during sprouting angiogenesis. We hypothesize that water flux across the cell membrane can lead to changes in tip cell volume, with water influx increasing the volume while outflow decreases. To quantify tip cell volume in the absence of Aquaporin function, we injected a plasmid encoding *kdrl:mEmerald* into 1-cell stage *aqp1a.1^rk28/rk28^;aqp8a.1^rk29/rk29^*embryos to label ECs in a mosaic manner. We also investigated the effects of increasing Aqp1a.1 expression on tip cell volume by injecting a plasmid encoding *Aqp1a.1-P2A-EGFP* into wildtype embryos. As control, wildtype embryos were injected with the plasmid encoding *kdrl:mEmerald*. mEmerald/EGFP-positive tip cells were imaged and their volume measured at 24 to 25 hpf from 3D reconstructions of labelled cells. Wildtype ECs are on average 964.6 ± 270.2 µm^3^ in size (Fig. 5A and D). In the absence of both Aqp1a.1 and Aqp8a.1 function (Fig. 5B), tip cell volume significantly decreased by 26% to 709.8 ± 293.5 µm^3^ (*p* = 0.015, Fig. 5D) when compared to wildtype tip cells. The overexpression of Aqp1a.1 increased tip cell volume by 36% to 1313 ± 527.4 µm^3^ when compared to wildtype tip cells (Fig. 5C and D, *p* = 0.0069) and by 85% when compared to *aqp1a.1^rk28/rk28^;aqp8a.1^rk29/rk29^*tip cells (*p* < 0.0001). These results therefore support water influx as the direction of water flow in tip cells during migration, and that this is mediated through Aqp1a.1 and Aqp8a.1.

**Figure 5.**
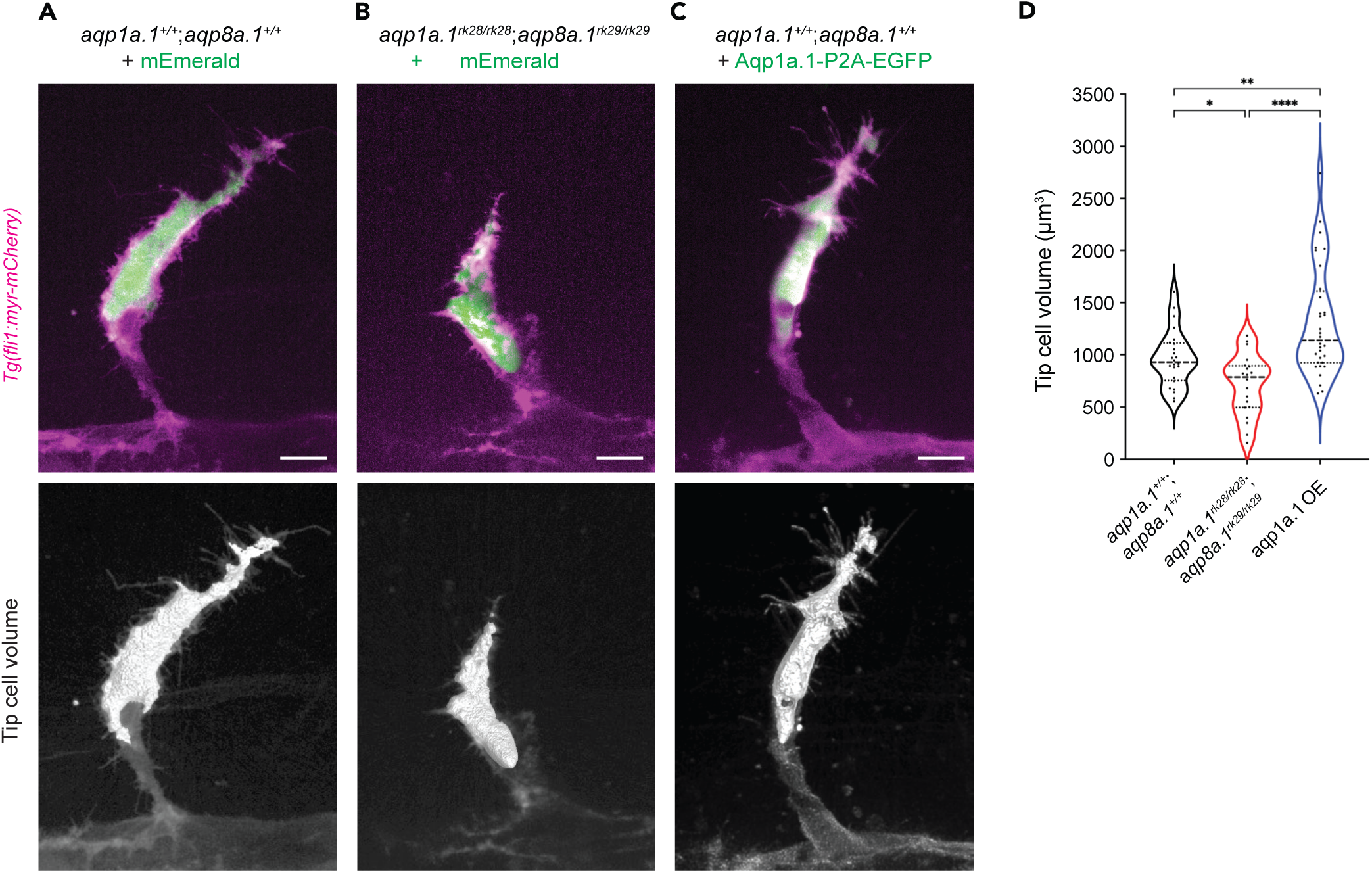
Aquaporin-mediated water influx increases tip cell volume. (**A-C**) Representative maximum intensity projection confocal z-stacks of tip cell and its 3D-volume rendering of 25 hpf wildtype (A), *aqp1a.1^rk28/rk28^;aqp8a.1^rk29/rk29^*(B) and wildtype embryos overexpressing Aqp1a.1 (C) labelled with cytosolic mEmerald or EGFP. Scale bar, 10 µm. (**D**) Quantification of tip cell volume in 25 hpf wild type (n = 24 cells) and *aqp1a.1^rk28/rk28^;aqp8a.1^rk29/rk29^* (n = 20 cells) and wildtype overexpressing Aqp1a.1 (n = 32 cells) embryos from 3 independent experiments. Statistical significance was determined by Brown-Forsythe and Welch ANOVA tests. *, *p* < 0.05, *\*\**, *p* < 0.01, and *\*\*\*\**, *p* < 0.0001.

In summary, our results demonstrate a role of endothelial Aquaporins in promoting water influx to increase endothelial tip cell volume and cytoplasmic hydrostatic pressure, as well as accelerate actin polymerization in filopodia (Fig. 4J).

### The anion channel, SWELL1, promotes EC migration and sprouting angiogenesis

As the direction of water flow is dictated by an osmotic gradient, with water flowing from low to high osmolarity, we next examined the function of ion channels in establishing an osmotic gradient. SWELL1, or LRRC8A, is a component of volume-regulated anion channel (VRAC) that mediates ions, especially chloride anions, and osmolyte efflux (Qiu et al., 2014; Voss et al., 2014). Recently, SWELL1 has been demonstrated to polarize at the trailing edge of migrating breast cancer cells to direct water efflux and confer confined migration direction (Zhang et al., 2022). In the zebrafish, scRNAseq data shows that the gene encoding SWELL1, *lrrc8aa*, is expressed in a subset of ECs, with 12.32% of ECs coexpressing *lrrc8aa* and *aqp1a.1* (Fig. 6A) and 10.16% coexpressing *lrrc8aa* and *aqp8a.1* (Fig. 6B). To determine whether *lrrc8aa* is expressed in endothelial tip cells, we performed RNAscope and confocal microscopy. Although *lrrc8aa* is expressed in surrounding somites and notochord (Fig. 6C), we could also detect its expression in endothelial tip cells (Fig. 6D and E), suggesting that SWELL1 can establish an osmotic gradient in the vicinity of the developing ISVs and within tip cells to direct water flow. To address the function of SWELL1-generated osmotic gradient, we disrupted its activity by treating zebrafish with 5µM DCPIB (Gunasekar et al., 2022) for 6 hours from 20 hpf and observed a similar impairment in ISV formation (Fig. 6F and G) as in *aqp1a.1^rk28/rk28^;aqp8a.1^rk29/rk29^* embryos (Fig. 3D and E). SWELL1 inhibition resulted in shorter ISVs formed at 26 to 27 hpf (Fig. 6H) due to impaired leading edge displacement of tip cells (Fig. 6I) and decreased tip cell velocity (Fig. 6J). Further analysis of tip cell morphology showed a decrease in the number (Fig. 6K) and length (Fig. 6L) of filopodia formed and compromised elongation of tip cells (Fig. 6M and N, Movie S7) in DCPIB-treated embryos. In summary, these results suggest that chloride efflux through SWELL1 establishes an osmotic gradient, either within ECs or between EC and the surrounding interstitial tissue, to direct water flow and thereby modulate EC migration and sprouting angiogenesis.

**Figure 6.**
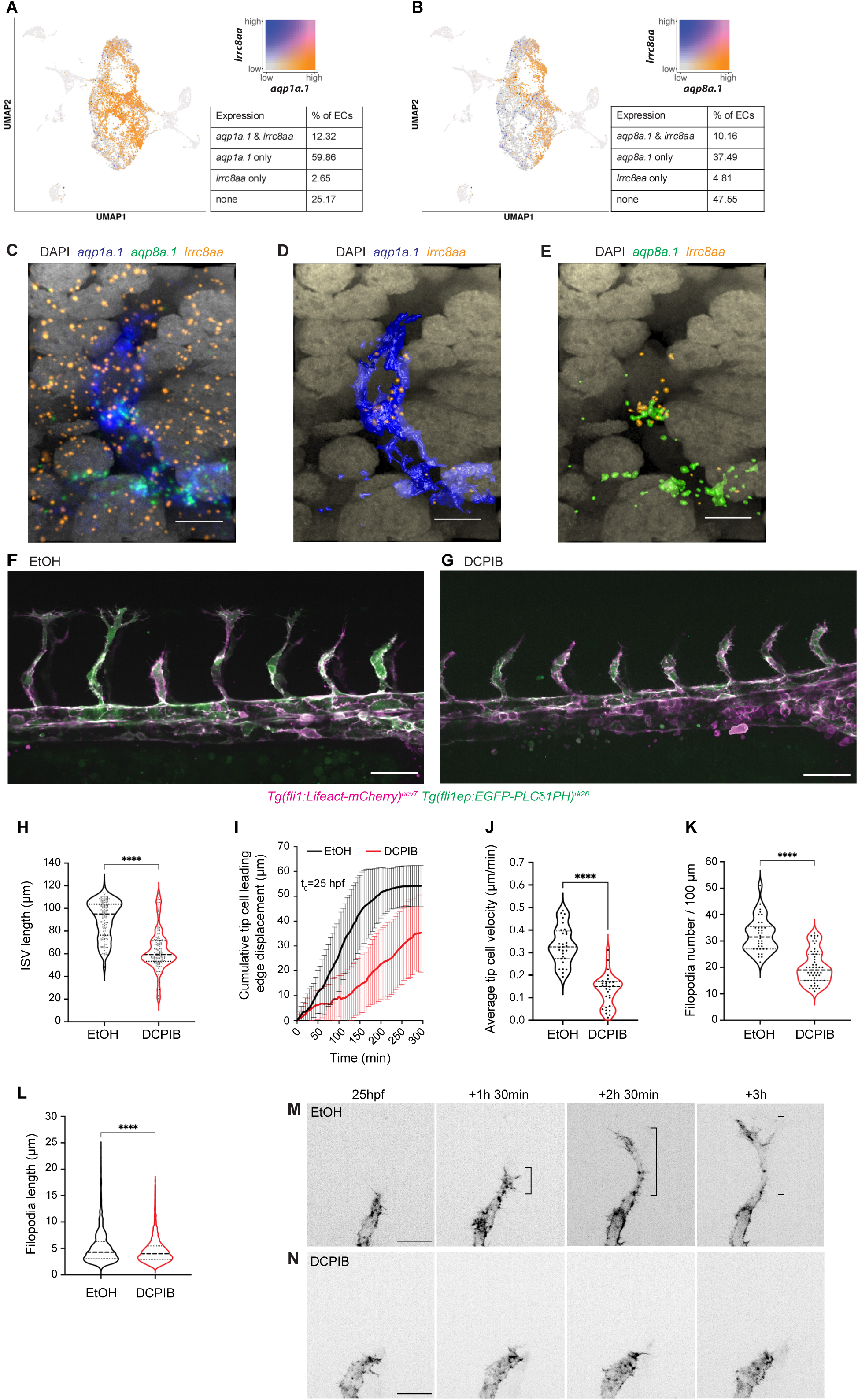
The chloride channel SWELL1 promotes EC migration and sprouting angiogenesis. (**A-B**) UMAP plots showing co-expression of *lrrc8aa* and *aqp1a.1* (A), and *lrrc8aa* and *aqp8a.1* (B) in endothelial cells from 24 hpf, 34 hpf and 3 dpf embryos. (**C-E**) Detection of *aqp1a.1*, *aqp8a.1* and *lrrc8aa* mRNA by RNAscope *in situ* hybridization in 22 hpf zebrafish embryo. Representative maximum intensity projection rendering of confocal z-stacks of tip and stalk cells (C). D and E show surface rendering of image C where only endothelial expression of *lrrc8aa* is shown after masking. Scale bar, 10 µm. (**F-G**) Representative maximum intensity projection confocal z-stacks of 26 hpf wildtype embryos treated with 0.05% EtOH (F) or 5 µM DCPIB (G) from 20 to 26 hpf (control, n = 16 embryos; DCPIB, n = 18 embryos; 2 independent experiments). Scale bar, 50 µm. (**H**) Quantification of ISV length in 26 hpf wildtype embryos treated with 0.05% EtOH or 5 µM DCPIB (control, n = 133 vessels from 16 embryos; DCPIB, 157 vessels from 18 embryos; 2 independent experiments). Statistical significance was determined with unpaired *t*-test; *\*\*\*\**, *p* < 0.0001. (**I**) Quantification of tip cell leading edge displacement of wildtype embryos treated with 0.05% EtOH (n = 29 cells from 7 embryos, 2 independent experiments) or 5 µM DCPIB (n = 29 cells from 8 embryos, 3 independent experiments) at 25 - 30 hpf. (**J**) Quantification of tip cell migration velocity in wildtype treated with 0.05% EtOH (n = 29 cells from 7 embryos, 2 independent experiments) or 5 µM DCPIB (n = 29 cells from 8 embryos, 3 independent experiments) at 25 - 30 hpf. Statistical significance was determined with unpaired *t*-test; *\*\*\*\**, *p* < 0.0001. (**K**) Quantification of filopodia number in tip cells of wildtype embryos treated with 0.05% EtOH (n = 34 vessels from 9 embryos, 3 independent experiments) or 5 µM DCPIB (n = 53 vessels from 16 embryos, 3 independent experiments). Statistical significance was determined with unpaired *t*-test; *\*\*\*\**, *p* < 0.0001. (**M-N**) Still images from representative time-lapse movies of migrating tip cells in wildtype embryos treated with 0.05% EtOH (M, n = 7) or 5 µM DCPIB (N, n = 8). Movies were taken from 25 to 30 hpf (n = 3 independent experiments). Bracket, formation of stable protrusions. Scale bar, 20 µm.

### Additive effects of actin polymerization and water influx in driving EC migration and sprouting angiogenesis

We have previously observed that ECs are still able to migrate and form ISVs after the inhibition of actin polymerization (Phng et al., 2013). In these experiments, embryos were treated with a low concentration of Latrunculin B (Lat. B), an inhibitor of actin polymerization, that resulted in the suppression of filopodia formation. Under such conditions, ECs can generate small membrane protrusions and migrate in a directed manner at a reduced speed. How ECs are still able to migrate after the inhibition of actin polymerization was unclear. As our current study demonstrates a role of water inflow and hydrostatic pressure as another mechanism of cell migration, we hypothesize that ECs employ water influx to migrate when actin polymerization is compromised and that the depletion of both water influx and actin polymerization will result in a greater inhibition of EC migration.

To test this hypothesis, we examined ISV development in embryos with decreased actin polymerization and water inflow by treating *aqp1a.1^rk28/rk28^;aqp8a.1^rk29/rk29^*embryos with Lat. B. At 28 hpf, ISVs of wildtype embryos treated with DMSO have reached the dorsal roof of the neural tube (Fig. 7A) and show an average length of 104.1 ± 13.47 µm (Fig. 7E). The treatment of wildtype embryos with 0.08 µg/ml Lat. B from 20 to 28 hpf (Fig. 7B) resulted in a significant decrease in ISV length (61.53 ± 23.27 µm) compared to control embryos (*p* < 0.0001, Fig. 7E). A similar decrease in ISV length was also observed in *aqp1a.1^rk28/rk28^;aqp8a.1^rk29/rk29^* embryos treated with DMSO (63.75 ± 30.24 µm, Fig. 7C and E). When *aqp1a.1^rk28/rk28^;aqp8a.1^rk29/rk29^* embryos were treated with 0.08µg/ml Lat. B from 20 to 28 hpf, there was a greater decrease in the length of ISV formed at 28 hpf (43.10 ± 24.42 µm, Fig. 7D and E) when compared to the inhibition of actin polymerization (*p* = 0.0028) or depletion of water influx (*p* < 0.0001) alone (Movie S8). These results therefore demonstrate additive functions of actin polymerization and water influx in driving EC migration during sprouting angiogenesis (Fig. 7F).

**Figure 7.**
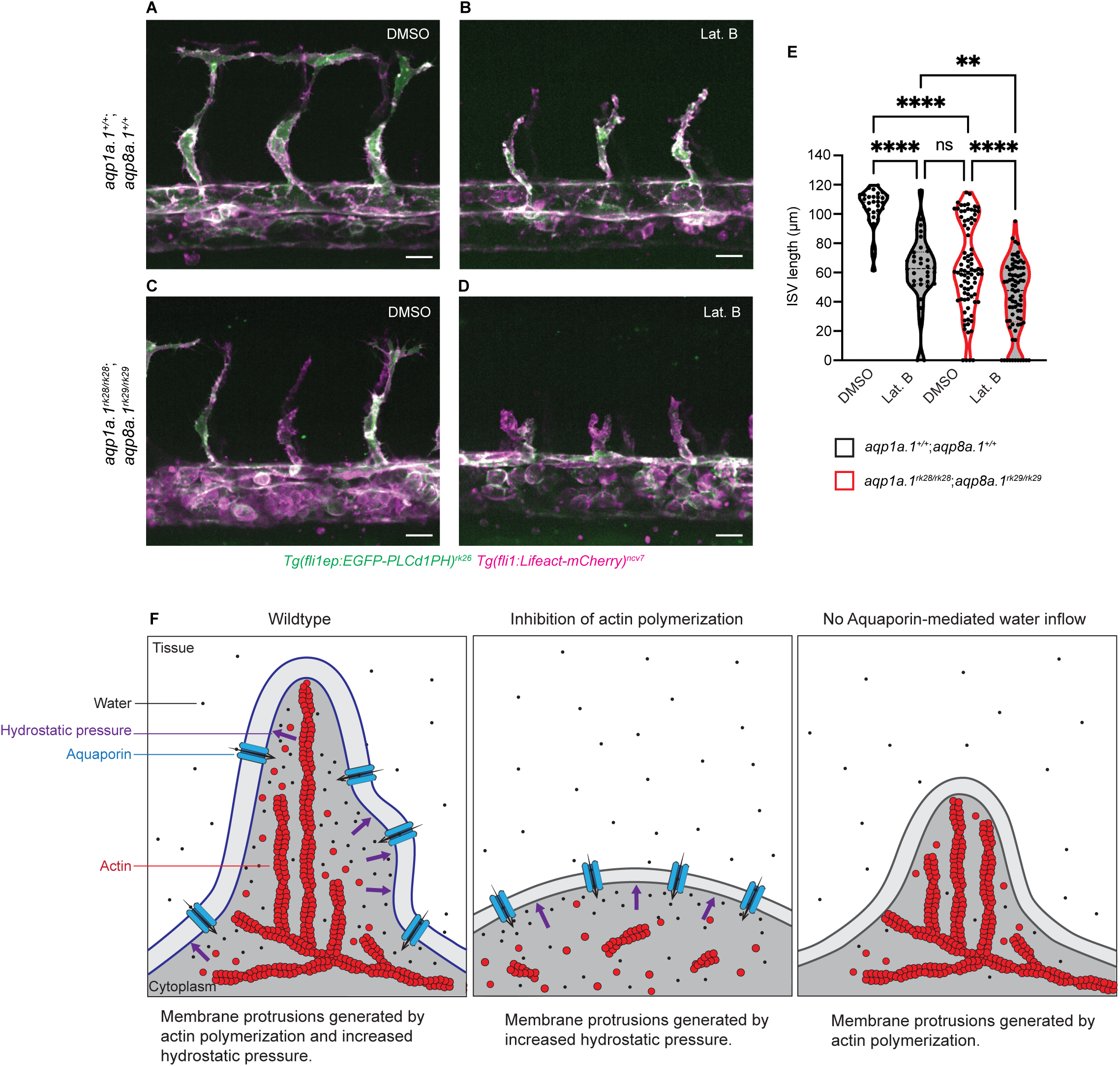
Additive function of actin polymerization and hydrostatic pressure in driving EC migration and sprouting angiogenesis. (**A-D**) Representative maximum intensity projection confocal z-stacks of 28 hpf wild type (A and B) and *aqp1a.1^rk28/rk28^;aqp8a.1^rk29/rk29^* (C and D) embryos treated with 0.1% DMSO (A and C) or 0.8 µg/ml Lat. B (B and D) from 20 to 28 hpf (for each condition: wild type, n = 10 embryos; *aqp1a.1^rk28/rk28^;aqp8a.1^rk29/rk29^*, n =12 embryos, 2 independent experiments). Scale bar, 20 µm. (**E**) Quantification of ISV length in 28 hpf wild type and *aqp1a.1^rk28/rk28^;aqp8a.1^rk29/rk29^*embryos treated with 0.1% DMSO or 0.8 µg/ml Lat. B (wild type: n = 60 vessels from 10 control embryos, n = 66 vessels from 10 Lat. B-treated embryos; *aqp1a.1^rk28^;aqp8a.1^rk29^*: n = 126 vessels from 12 control embryos, n = 112 vessels from 11 Lat. B-treated embryos, 2 independent experiments). Statistical significance was determined by Brown-Forsythe ANOVA test with Sidak’s multiple comparison test; ns, *p,*>,0.05, *^**,^p* < 0.01, and *^****,^ p* < 0.0001. (**F**) Model of endothelial tip cell migration. Tip cells generate membrane protrusions by actin polymerization and increased hydrostatic pressure via Aquaporin-mediated water inflow. In the absence of actin polymerization and presence of Aquaporins, hydrostatic pressure deforms membranes to generate membrane protrusions. When Aquaporin function is lost, membrane protrusions is generated by actin polymerization. Fewer membrane protrusions are formed when only one mechanism is utilized, resulting in slower EC migration. See also Movie S8.

## DISCUSSION

In this study, we demonstrate that endothelial tip cells utilize two mechanisms of migration concurrently – actin polymerization and hydrostatic pressure - during sprouting angiogenesis in the zebrafish. The increase in hydrostatic pressure in endothelial tip cells is generated by Aqp1a.1- and Aqp8a.1-mediated water inflow. By performing detailed expression analyses, we showed that *aqp1a.1* and *aqp8a.1* are differentially expressed in newly formed vascular sprouts. *Aqp1a.1* is expressed earlier in the DA than *aqp8a.1* during development and in ECs destined to be tip cells. In newly formed ISVs, endothelial tip cells express higher *aqp1a.1* mRNA levels while stalk cells express higher *aqp8a.1* mRNA levels. In zebrafish harbouring mutations in both *aqp1a.1* and *aqp8a.1*, there is delayed tip cell protrusion from the DA, decreased tip cell migration and increased occurrence of truncated ISVs at 3 dpf. We further discovered that *aqp1a.1* and *aqp8a.1* expression is upregulated by VEGFR2 activity, suggesting that VEGFA-VEGFR2 signalling employs Aquaporin function as one mechanism to drive sprouting angiogenesis.

Several lines of evidence implicate Aquaporins in facilitating water flow into endothelial tip cells during sprouting angiogenesis. First, timelapse imaging showed polarized localization of Aqp1a.1 and Aqp8a.1 proteins at the leading edge of migrating tip cells, suggesting that water flow occurs locally at the migrating front of the cell. Secondly, single cell analyses revealed Aqp1a.1- and Aqp8a.1-deficient tip cells have reduced cell volume, impaired expansion of cell membranes at the leading edge and decreased generation of stable cell protrusions. These observations implicate water flow into tip cells, leading to increased cell volume. This is confirmed by the overexpression of Aqp1a.1 in tip cells that resulted in increased cell volume. Due to the incompressible property of water within an enclosed compartment such as the cell, water influx will lead to elevation in hydrostatic pressure. At the leading edge of tip cells, increased hydrostatic pressure would expand and deform membranes to generate protrusions. Additionally, water influx can reduce cytoplasmic viscosity and increase spacing between the plasma membrane and the underlying actin cytoskeleton, promoting actin monomer diffusion, actin polymerization and the generation of membrane protrusions to facilitate cell migration (Boer et al., 2023; Loitto et al., 2009).

A key finding in this study is the demonstration that endothelial tip cells employ hydrostatic pressure as a second mechanism of cell migration *in vivo*. In a previous study, we discovered that ECs continue to migrate when actin polymerization is inhibited and in the absence of filopodia (Phng et al., 2013). Here, we show that hydrostatic pressure can generate force for EC migration in the absence of actin-based force generation. When both mechanisms are perturbed, there is a greater impairment in EC migration (Fig. 7D and E). This finding highlights the importance of hydrostatic pressure as an additional mechanism that reinforces actin-based cell shape changes and motility to ensure robust migration and formation of new blood vessels in complex three-dimensional environments. Such dual mode migration is advantageous when blood vessels develop in physically confined spaces such as in the zebrafish trunk, where ECs must emerge from the DA and migrate between somite boundaries and over tissues such as the notochord and neural tube to form ISVs. Such dependence on water flow in cell migration under confined microenvironment has been reported in human MDA-MB-231 breast cancer cells and mouse S180 sarcoma cells, which show cell migration persistence in narrow channels after inhibition of actin polymerization or myosin II-mediated contractility (Stroka et al., 2014). In this context, AQP5 mediates water flow and the decrease or increase in AQP5 expression suppresses or enhances, respectively, cancer cell migration (Chae et al., 2008; Jung et al., 2011; Stroka et al., 2014).

Our findings on the endothelial function of zebrafish Aqp1a.1, which is homologous to mammalian AQP1, corroborate previous studies demonstrating a function of AQP1 in promoting cell migration. In tumour bearing AQP1-null mice, there is reduced tumour angiogenesis and AQP1-deficient ECs display reduced migration in an *in vitro* wound healing assay (Maltaneri et al., 2020; Saadoun et al., 2005). In chick, AQP1 expression levels affect neural crest cell migration speed and direction by regulating filopodia length and stability (McLennan et al., 2019). The regulation of membrane protrusions has also been demonstrated for other Aquaporin proteins. For example, AQP9 weakens membrane-cytoskeleton anchorage and promote formation of membrane protrusions such as filopodia and blebs (Karlsson et al., 2013; Loitto et al., 2007). We also demonstrate for the first time a role of Aqp8a.1 in promoting EC migration and sprouting angiogenesis in the zebrafish. However, its mammalian ortholog, AQP8, was shown to inhibit colorectal cancer cell lines *in vitro* (Wu et al., 2017), suggesting cell- and context-dependent function of AQP8 in the regulation of cell migration. The role of Aqp1a.1 and Aqp8a.1 has also been recently investigated in the zebrafish. However, the authors did not report defects in EC migration but instead observed altered ISV diameter in Aqp1a.1 and Aqp8a.1 knockout and overexpression experiments (Chen et al., 2024). While we corroborate a decrease in the diameter of aISVs and vISVs in *aqp1a.1^rk28/rk28^*and *aqp1a.1^rk28/rk28^;aqp8a.1^rk29/rk29^* zebrafish, we observed a slight increase in the diameter in *aqp8a.1^rk29/rk29^*zebrafish at 2 dpf (Fig. S11A). Furthermore, unlike described in Chen et al, we could not detect a difference in ISV diameter between control and Aqp1a.1- or Aqp8a.1-overexpressing ECs (Fig. S11B). The disparity in phenotypes observed between our groups are likely a result of differences in the *aqp1a.1* and *aqp8a.1* mutations and the level of Aqp1a.1 or Aqp8a. overexpression generated in our respective zebrafish lines.

A central determinant of water movement is the osmotic gradient across the membrane, with water flowing in the direction of higher osmotic concentration to build up hydrostatic pressure. Given a crucial role of water flow in regulating cell volume, shape and migration, it is also important to understand the function and distribution of ion channels, exchangers and transporters and water channels that generate an osmotic gradient across the cell. This is reflected in a growing number of studies demonstrating a role of ion transporters in cell migration. In neutrophils, the increase in cell volume and potentiation of cell migration depend on the sodium-proton exchanger 1 (NHE1) and the chloride-bicarbonate exchanger 2 (AE2) (Nagy et al., 2023). In T cells, chemokine-induced migration depends on ion influx via SLC12A1, and water influx via AQP3 (Boer et al., 2023). In tumour cells migrating in narrow channels, the establishment of a polarized distribution of Na^+^/H^+^ pumps and Aquaporin proteins in cell membranes is required for net inflow of ions and water at the leading edge and net outflow at the trailing edge (Stroka et al., 2014). Although our study identifies a function of Aqp1a.1 and Aqp8a.1 in facilitating water inflow, it remains unknown which ion channels and transporters control ion flux across the cell membrane to modulate water flow in tip cells. Results from our study suggest a role of SWELL1 in generating an osmotic gradient that controls EC migration, since its inhibition resulted in reduced membrane protrusions, slower tip cell migration and defective sprouting angiogenesis. Due to the limitations of our experimental approach, we do not know whether the effect of inhibiting SWELL1 function is cell autonomous or a consequence of altered chloride flux in the neighbouring cells that would alter tissue osmolarity. It also remains to be clarified where SWELL1 is localized within migrating ECs, how it regulates the osmotic gradient within the EC (cellular) and between the EC and the surrounding interstitial tissue (extracellular) to control the direction of water flow, and whether other ion channels also work in concert to modulate overall cellular and extracellular osmolarity to direct EC migration.

We have also observed enrichment of Aqp1a.1 and Aqp8a.1 proteins in specific compartments of ECs, at the leading edge of migrating endothelial tip cells as well as apical membranes during lumen formation (not shown). This leads to the question of how Aquaporin localization is regulated as this will determine the site of increased water inflow and hydrostatic pressure. It has been shown that rapid changes in subcellular localization of mammalian Aquaporins upon stimulation occurs mainly via trafficking to and from the plasma membrane (Markou et al., 2022) to modulate the amount of protein in the plasma membrane. The regulation of AQP subcellular relocalization via calmodulin- and/or phosphorylation-dependent mechanisms has been implicated for AQP0-5 and AQP7-9 (Markou et al., 2022). For example, rapid translocation of AQP1 to plasma membrane upon a hypotonic stimulus is dependent on calmodulin activation and phosphorylation by protein kinases C (PKC) (Conner et al., 2012) and subcellular localization of AQP8 in hepatocytes is also regulated by PKA and PI3K signalling (Gradilone et al., 2005, 2003). Further work is needed to elucidate the mechanism of Aqp1a.1 and Aqp8a.1 protein distribution in ECs.

In this study, we have assumed that Aqp1a.1 and Aqp8a.1 regulate EC behaviours through their ability to control water permeation based on previous studies demonstrating that water molecules flow through mammalian AQP1 (Preston et al., 1992; Zeidel et al., 1992) and AQP8 (Liu et al., 2006; Ma et al., 1997) as well as through zebrafish orthologs Aqp1a.1 and Aqp8a.1 (Tingaud-Sequeira et al., 2010). However, AQP1 and AQP8 are not water-specific channels. AQP1 can conduct unpolar gases such as carbon dioxide (Nakhoul et al., 1998; Prasad et al., 1998; Wang et al., 2007) and nitric oxide (Herrera et al., 2006; Herrera and Garvin, 2007), and in zebrafish, Aqp1a.1 also mediates the transfer for small gaseous molecules such as carbon dioxide and ammonia (Talbot et al., 2015). AQP8 is permeable to ammonia (Jahn et al., 2004; Liu et al., 2006) and is present in the inner mitochondrial membrane (Calamita et al., 2005), where it is suggested to mediate ammonia transport rather than water fluxes (Soria et al., 2010). Nevertheless, because water movement flows up an osmotic gradient, increased entry of ions into the cell through Aquaporins will also trigger water inflow.

In summary, our study highlights the role of water influx and hydrostatic pressure as another force-generating mechanism utilized by cells to build tissues during animal development. We demonstrate that endothelial tip cells employ both actin polymerization and hydrostatic pressure for robust sprouting angiogenesis. As morphogenetic events are governed by a coordination of cell shape changes, cell migration and rearrangements, we envision that the shaping and formation of other tissues will also depend on Aquaporin function and changes in hydrostatic pressure.

## METHODS

### Zebrafish maintenance and stocks

Zebrafish (*Danio rerio*) were raised and staged according to established protocols (Kimmel et al., 1995). Transgenic lines used in this work are *Tg(kdrl:EGFP)^s843^* (Jin et al., 2005), *Tg(fli1a:H2B-EGFP)^ncv69^* [re-named in ZFIN as *Tg(fli1:zgc:114046-EGFP)*] (Ando et al., 2019), *Tg(fli1:Lifeact-mCherry)^ncv7^*(Wakayama et al., 2015), *Tg(fli1:myr-mCherry)^ncv1^* (Fukuhara et al., 2014), *Tg(fli1ep:EGFP-PLC1δPH)^rk26^* (Kondrychyn et al., 2020), *Tg(fli1:h2bc1-mCherry)^ncv31^* (Nakajima et al., 2017), *Tg(kdrl:ras-mCherry)^s916^* (Hogan et al., 2009), *Tg(kdrl:nls-EGFP)^ubs1^;(fli1:Has.B4GALT1-mCherry)^bns9^* (Kwon et al., 2016), *Tg(fli1ep:GAL4FF)^ubs2^*(Herwig et al., 2011). Zebrafish were maintained on a 12-hour light/12-hour dark cycle, and fertilized eggs were collected and raised in E3 medium at 28^0^C. To inhibit pigmentation in embryos older than 24 hpf, 0.003% *N*-Phenylthiourea (Sigma-Aldrich) in E3 medium was used. All animal experiments were approved by the Institutional Animal Care and Use Committee at RIKEN Kobe Branch (IACUC).

### Plasmid construction and transgenesis

Plasmid *mEmerald-N1* was a gift from Michael Davidson (Addgene plasmid #53976), plasmids *pMDS6* containing terminal *cis*-required sequences of the *Ds* transposon from maize and *pAc-SP6* containing an ORF of *Ac* transposase were a gift from Sergei Parinov. Other plasmids used in this study were generated by using In-Fusion HD Cloning kit (Takara Bio Inc.). A detailed information regarding plasmids and primers used in this study can be found in Supplementary Tables 1 and 2, respectively. *Tg(fli1ep:aqp1a.1-mEmerald)^rk30^* and *Tg(fli1ep:aqp8a.1-mEmerald)^rk31^* transgenic zebrafish were generated by injecting *Tol2*-based *fli1ep:aqp1a.1-mEmerald* or *fli1ep:aqp8a.1-mEmerald* plasmids (10 ng/μl), respectively, along with *Tol2 transposase* mRNA (100 ng/μl) into one-cell stage AB embryos. *Tol2* transposase mRNA was transcribed from Not*I*-linearized *pCS-TP* plasmid (a gift from Koichi Kawakami, National Institute of Genetics, Japan) using the mMESSAGE mMACHINE SP6 kit (Invitrogen). Injected embryos were raised to adult and screened for founders.

### Cloning of *aqp1a.1* and *aqp8a.1* genes

Total RNA was isolated from 1-day old zebrafish embryos with TRI Reagent using Direct-zol^TM^ RNA MicroPrep kit (Zymo Research) according to the manufacturer’s protocol. The first-strand cDNA was synthesized from 1 µg of a total RNA by oligo(dT) priming using the SuperScript III First-Strand synthesis system (Invitrogen) according to the manufacturer’s protocol. Amplification of cDNAs was performed using a high-fidelity KOD-Plus-Neo DNA polymerase (Toyobo, Japan) and resulting PCR products were cloned using NEB^®^ PCR Cloning kit (New England BioLabs). Positive clones and plasmids were verified by DNA sequencing.

### Mosaic expression of DNA constructs

To avoid unnecessary *Tol2* re-transposition from the donor site in transgenic zebrafish lines generated using *Tol2* transposon (Kondrychyn et al., 2009), *Ac*/*Ds* transposon system from maize (Emelyanov et al., 2006) was used for mosaic expression of DNA constructs. *Ds*-based plasmids (10 pg) were co-injected with 100 pg of *Ac* transposase mRNA, transcribed from Bam*HI*-linearized *pAc-SP6* plasmid using the mMESSAGE mMACHINE SP6 kit (Invitrogen), into one-cell stage zebrafish wildtype or *aquaporin* mutant embryos. Embryos were analyzed at 25-26 hpf after injections.

### *In vivo* cell volume analysis

Single-cell labelling of ECs in ISVs was achieved by mosaic expression of *pDs-kdrl:mEmerald* plasmid in wild type or *aqp1a.1^rk28/rk28^;aqp8a.1^rk29/rk29^*mutant *Tg(fli1:Lifeact-mCherry)^ncv7^* transgenic embryos. To visualize single ECs with Aqp1a.1 overexpression, we injected *pDs-6xUAS:aqp1a.1-P2A-EGFP-T2A-mKate2-CAAX* plasmid into *Tg(fli1:GAL4FF)^ubs2^; Tg(fli1:myr-mCherry)^ncv1^* embryos. Embryos were imaged at 25-26 hpf with an Olympus UPLSAPO 60x/NA 1.2 water immersion objective with optical Z planes interval of 0.26 μm. Cell volume was measured with Huygens Essential 22.10 software (Scientific Volume Imaging B.V.) using object analyzer tool with manual adjustment of threshold.

### Generation of zebrafish *aqp1a.1* and *aqp8a.1* mutants

Aquaporins mutants were generated by CRISPR/Cas9-mediated mutagenesis. Guide RNAs (gRNAs) targeting the first exons of *aqp1a.1* and *aqp8a.1* (Fig. S4 and S5) were generated by using cloning-independent protocol (Gagnon et al., 2014). Briefly, to generate templates for gRNA transcription, a 60-base oligodeoxynucleotides containing the SP6 promoter sequence, the gene-specific sequence (*aqp1a.1*: 5’-GACAGCTGGCCAGCAGACCC-3’; *aqp8a.1*: 5’-GATGTCTCCCCCATCGCCCG-3’) and 23-base overlap region was annealed to an 80-base constant oligodeoxynucleotide encoding the reverse-complement of the tracrRNA tail. The ssDNA overhangs were filled-in with T4 DNA polymerase (New England BioLabs) to generate dsDNA. The gRNAs were *in vitro* transcribed using MEGAScript SP6 kit (Invitrogen), DNase treated and purified with RNA Clean and Concentrator kit (Zymo Research). The gRNAs were quality-checked by running 1 µg of the product on a 2% TBE-agarose gel. Cas9/gRNA RNP complex was assembled just prior to injection and after 5 min incubation at room temperature 200 pg of Cas9 protein (Invitrogen) and 200 pg of gRNAs (100 pg of each, *aqp1a.1* gRNA and *aqp8a.1* gRNA) were co-injected into one-cell stage *Tg(fli1:myr-mCherry)^ncv1^* embryos. Such dose produced over 70% embryos survival post-injection showed CRISPR/Cas9-induced somatic *aqp1a.1* and *aqp8a.1* gene mutations. F_0_ founders were identified by outcrossing CRISPR/Cas9-injected fish with wild type fish and screening the offspring for mutations at 2 dpf using Sanger sequencing. We found one F_0_ fish with germline transmitted mutations in both *aqp1a.1* and *aqp8a.1* genes and outcrossed it with *Tg(fli1:myr-mCherry)^ncv1^*and *Tg(fli1a:H2B-EGFP)^ncv69^* transgenic fish to establish F_1_ generation of heterozygotes. A 55 F_1_ fish were genotyped to identify 4 *aqp1a.1^+/rk28^*, *11 aqp8a.1^+/rk29^* and 12 *aqp1a.1^+/rk28^;aqp8a.1^+/rk29^* heterozygote fish. Subsequently, F_1_ heterozygote fish were in-crossed to establish a single and double homozygote fish.

### DNA isolation and genotyping

Genomic DNA was isolated from either embryos or fin clips using HotSHOT method (Meeker et al., 2007). To identify genomic lesions, the following primers were used: aqp1a1-fwd (5’-CGCCTCCAGATTCATTAGCAGGA-3’) and aqp1a1-rev (5’-GTAAGTGAACTGCTGCCAGTGA-3’) to amplify a 550 bp fragment, and aqp8a1-fwd (5-GGATCAATTGAGTTGCATAACAGAC-3’) and aqp8a1-rev (5’-CTGTAATGTAGACTTGTAAAGTGGA-3’) to amplify a 638 bp fragment. Mutations were assessed by direct sequencing of purified PCR products.

### Quantitative real-time PCR

Total RNA from whole embryos was isolated with TRI reagent using Direct-zol RNA MicroPrep kit (Zymo Research) according to the manufacturer’s protocol, including on-column DNA digestion. RNA was quantified using a NanoDrop 1000 spectrophotometer (ThermoFisher Scientific, USA). cDNA was synthesized from 300 ng of purified RNA using LunaScript RT SuperMix kit (New England BioLabs) according to the manufacturer’s protocol. Amplification of target cDNA was performed in technical triplicate of three biological replicates using the SYBR green methods. Each qPCR reaction mixture contained 5 µL 2x Luna Universal qPCR master mix (New England BioLabs), 1 µL cDNA (2-fold dilution), 0.2 µL antarctic thermolabile UDG (New England BioLabs) and 250 nM each primer to a final volume of 10 µL. Reactions were run in 384-well plates (Applied Biosystems) using the QuantStudio 5 Real-Time PCR system (Applied Biosystems) with the following thermal cycling conditions: initial UDG treatment at 25^0^C for 2 minutes, UDG inactivation at 50^0^C for 5 min and denaturation at 95^0^C for 1 minutes, followed by 45 cycles of 15 sec at 95^0^C, 30 sec at 60^0^C with a plate read at the end of the extension step. Control reactions included a no-template control (NTC) and a no-reverse transcriptase control (NRT). Dissociation analysis of the PCR products was performed by running a gradient from 60 to 95^0^C to confirm the presence of a single PCR product. Amplification data was analyzed using QuantStudio Design and Analysis software v1.5.3 (Applied Biosystems) and relative fold change was calculated using Pfaffl method (Pfaffl, 2001) and normalized to *gapdh* expression (as an internal control). Primers are listed in Supplementary Table 2. Two embryos were used in each biological replicate to generate an average value that was used to calculate the final mean ± SD from two or three independent experiments.

### RNA *in situ* hybridization

#### RNAscope *in situ* hybridization

RNAscope^®^ *in situ* hybridization was conducted by using RNAscope Multiplex Fluorescent Reagent kit v2 (Advanced Cell Diagnostics). We adapted manufacture’s protocol designed for samples mounted on slides and protocol developed for whole-mount embryo samples (Gross-Thebing et al., 2014). For each experimental point, 5 embryos were processed in one 1.5 ml Eppendorf tube. Briefly, 20 and 22 hpf old embryos were manually dechorionated and fixed in freshly prepared 4% methanol-free PFA in PBS for 1 hour at room temperature. After fixation, embryos were washed three times in PBS containing 0.01% Tween-20 (PBST), dehydrated stepwise in 5-min washes in a series of increasing methanol concentrations (25%, 50%, 75%, 100%) in PBS and then stored in 100% methanol at -20^0^C before use them for hybridization. The methanol-stored embryos were incubated with 5% H_2_O_2_ in methanol for 20 min at room temperature and then rehydrated stepwise in 5-min washes in series of decreasing methanol concentrations (75%, 50%, 25%) in PBS, followed by three times 5-min washes in PBST. Embryos were permeabilized in 2 drops of RNAscope^®^ Protease III for 15 min, rinsed three times with PBST and hybridized with RNAscope target probes overnight at 50^0^C in water bath. After hybridization probes were recovered and embryos were washed three times with 0.2x SSCT [0.01% Tween-20 in SSC] at room temperature, re-fixed with 4% PFA for 10 min, and washed three times with 0.2x SSCT. For RNA detection, the embryos were first hybridized with three different amplifier solution for 30 min (Amp1, and Amp2) and 15 min (Amp3) in a water bath at 40^0^C. After each hybridization step, the embryos were washed three times with 0.2x SSCT for 10 min at room temperature. To develop signal of each probe, the embryos were sequentially incubated in a water bath at 40^0^C with: (i) the horseradish peroxidase (HRP) for 15 min, (ii) TSA fluorophore for 30 min and (iii) the HRP blocker for 15 min. After each incubation step, the embryos were washed three times with 0.2x SSCT for 10 min at room temperature. HRP step is linked to a probe channel, we first developed C1 probe, *Dr-aqp1a.1* (Advanced Cell Diagnostics) using HRP-C1 and TSA Vivid Fluorophore 520 (Tocris, dilution 1:1500), then C2 probe, *Dr-lrrc8aa* (Advance Cell Diagnostics) using HRP-C2 and TSA Vivid Fluorophore 570 (Tocris, dilution 1:1500), and finally C3 probe, *Dr-aqp8a.1* (Advanced Cell Diagnostics) using HRP-C3 and TSA Vivid Fluorophore 650 (Tocris, dilution 1:1500). Nuclei were counterstained with DAPI ready-to-use solution overnight at 4^0^C in the dark. Prior to imaging embryos were rinsed in PBST and kept in 70% glycerol in PBS at 4^0^C in the dark. Images were acquired using Olympus FV3000 confocal microscope and an Olympus UPlanXApo 60x/NA 1.42 oil immersion objective.

#### Chromogenic *in situ* hybridization

Chromogenic *in situ* hybridization was conducted according to standard protocol (Thisse and Thisse, 2007) with minor modifications. For each experimental point, 10 embryos were processed in one 1.5 ml Eppendorf tube. Briefly, 30, 48, 72 and 96 hpf old embryos were manually dechorionated and fixed in freshly prepared 4% methanol-free PFA in PBS (pH 7.4) at 4^0^C overnight. After fixation embryos were washed three times in PBS containing 0.1% Tween-20 (PBST), dehydrated stepwise in 5-min washes in a series of increasing methanol concentrations (25%, 50%, 75%, 100%) in PBS and then stored in 100% methanol at -20^0^C before use them for hybridization. The methanol-stored embryos were rehydrated stepwise in 5-min washes in series of decreasing methanol concentrations (75%, 50%, 25%) in PBS followed by three times 5-min washes in PBST, and permeabilized 20 min with 10 µg/mL proteinase K (ThermoFisher Scientific). Hybridization was carried out in buffer [50% formamide, 5x SSC, 50 g/mL heparin, 500 g/mL tRNA (Roche) and 0.1% Tween-20] containing 5% dextran sulfate (MW=500 kDa) at 69^0^C overnight. After stringency wash, the specimens were blocked with 2% blocking reagent (Roche) in maleic acid buffer [100 mM maleic acid, 150 mM NaCl, 50 mM MgCl_2_ and 0.1% Tween-20, pH 7.5] and incubated overnight with anti-digoxigenin antibody, conjugated with alkaline phosphatase (dilution 1:5000, Roche) at 4^0^C. For detection of alkaline phosphatase, specimens were incubated in staining buffer [50 mM Tris-HCl, 50 mM NaCl, 25 mM MgCl_2_, 2% Polyvinyl alcohol and 0.1% Tween-20, pH 9.5] containing 375 µg/mL nitro-blue tetrazolium chloride (Roche) and 175 µg/mL 5-bromo-4-chloro-3’-indolyl-phosphate (Roche). Embryos were cleared in 70% glycerol overnight and imaged using Leica M205FA microscope.

#### RNA probe synthesis

The cDNA-containing vectors were linearized with appropriate restriction enzymes (detailed maps of vectors will be provided upon request) and used as a template for RNA probe synthesis. Sense and antisense RNA probes were synthesized using MEGAscript SP6 or T7 kits (Invitrogen) and digoxigenin (DIG)-labeled rNTPs (Roche) according to the manufacturer’s protocol and purified using RNA Clean and Concentrator kit (Zymo Research). The following sense and antisense DIG-labeled riboprobes were generated: (i) *aqp1a.1*, 1.1 kb long probe comprised 121 nt of 5’UTR, 783 nt of the open reading frame (ORF) and 207 nt of 3’UTR; (ii) *aqp8a.1*, 0.8 kb long probe comprised 17 nt of 5’UTR, 783 nt of ORF and 30 nt of 3’UTR. Sense probes showed no specific or unspecific staining (data not shown).

#### Image processing

Chromogenic and RNAscope *in situ* hybridization images were processed using ImageJ software (NIH, version 2.9.0/1.53t) with brightness and contrast adjustments. Z-stacks of fluorescent images are presented as maximum intensity projections.

### Analysis of mRNA expression in tip and stalk cells

RNAscope processed embryos were mounted on slide and imaged using Olympus FV3000 confocal microscope and Olympus UPlanXApo 60x/NA 1.42 oil immersion objective. Maximum intensity projection of z-stacks was used to determine the level of cellular fluorescence in ImageJ. The tip and stalk cells were selected using freehand selection tool, “mean grey value”, “area” and “integrated density” in each ROI were measured first for channel 1 (aqp1a.1). A region next to cell that has no fluorescence was selected as background and measured. The step was repeated for channel 2 (aqp8a.1) on the same ROIs. Two parameters were calculated for each channel, (i) the corrected total cell fluorescence, CTCF=Integrated Density – (Area of selected cell x Mean fluorescence of background) and (ii) ratio, R=CTCF of tip cell/CTCF of stalk cell. We consider that if R <0.9, expression is lower in tip cell, if R=0.9-1.1, expression is equal in tip and stalk cells, if R>1.1, expression is higher in tip cell.

### Imaging

For live confocal imaging, embryos were mounted in 0.8% low-melt agarose (Bio-Rad) in E3 medium containing 0.16 mg/mL Tricaine (Sigma-Aldrich) and 0.003% phenylthiourea (Sigma-Aldrich) in glass bottom 35-mm dishes (MatTek). Confocal Z-stacks were acquired using an inverted Olympus IX83/Yokogawa CSU-W1 spinning disc confocal microscope equipped with a Zyla 4.2 CMOS camera (Andor) and Olympus UPLSAPO 40x/NA 1.25 or 30x/NA 1.05 silicone oil immersion objectives. Bright-field images were acquired on Leica M205FA microscope. Images were processed using ImageJ software (NIH, version 2.9.0/1.53t).

### Analysis of cell migration

*aqp1a.1^+/rk28^;aqp8a.1^+/rk29^* and *aqp1a.1^rk28/rk28^;aqp8a.1^rk29/rk29^*embryos on *Tg(fli1a:H2B-EGFP)^ncv69^*;*(fli1:Lifeact-mCherry)^ncv7^* double transgenic background were imaged from 21 to 30 hpf with time interval 6 min using Olympus UPLSAPO 30x/NA 1.05 silicon oil immersion objective. We tried to mount both heterozygote and mutant embryos on the same dish (1 het plus 2 mut, or 2 het plus 2 mut) to image embryos on the same experimental conditions. Acquired time-lapse images were registered using HyperStackReg plugin in ImageJ, tip cell and nuclei were tracked using Manual Tracking plugin and velocity was calculated.

### Analysis of grow rate of actin bundles in filopodia

Wild type and *aqp1a.1^rk28/rk28^;aqp8a.1^rk29/rk29^*embryos on *Tg(fli1ep:EGFP-PLC1δPH)^rk26^;(fli1:Lifeact-mCherry)^ncv7^* double transgenic background were imaged starting from 25 hpf for 10 min with time interval 30 sec using Olympus UPLSAPO 60x/NA 1.2 water immersion objective. Actin bundles growth was tracked using Manual Tracking plugin and velocity was calculated.

### Analysis of filopodia number and length

Wild type, *aqp1a.1^+/rk28^;aqp8a.1^+/rk29^* and *aqp1a.1^rk28/rk28^;aqp8a.1^rk29/rk29^* embryos on *Tg(fli1ep:EGFP-PLC1δPH)^rk26^;(fli1:Lifeact-mCherry)^ncv7^*double transgenic background were imaged starting from 24 hpf for 30 min with time interval of 50 sec using Olympus UPLSAPO 60x/NA 1.2 water immersion objective. In a single image we counted filopodia number per vessel, measured membrane length and calculated filopodia number.

### Assessment of tip cell polarization

*Tg(kdrl:nls-EGFP)^ubs1^;(fli1:Has.B4GALT1-mCherry)^bns9Tg^*wildtype and *aqp8a.1^rk29/rk29^* embryos were imaged at 24-26 hpf using an Olympus UPLSAPO 40x/NA 1.25 silicon oil immersion objective. To analyze endothelial tip cell polarity in *aqp1a.1^rk28/rk29^* embryos, Golgi apparatus and nuclei were labelled by mosaic expression of *pDs-fli1ep:nls-EGFP-P2A-mKate2-GM130* plasmid. Images were analyzed using Fiji (NIH). To assess polarity, the nucleus was fit into an ellipsoid shape and the angle between the primary axis of the ellipse and the center of the Golgi was measured (Fig. S10D). Polarization of ECs is defined as follows: (1) front (polarized), if the Golgi is located within -30^0^ ∼ +30^0^; (2) none, if the Golgi is located within +30^0^ ∼ +150^0^ or -30^0^ ∼ -150^0^; or (3) rear, if the Golgi is located on the downstream side of the nucleus and angle is -150^0^ ∼ +150^0^. Tip cells in ISVs no. 5-15 were analyzed.

### Quantification of EC number and ISV diameter

Measurements were performed in wildtype, *aqp1a.1^rk28/rk28^, aqp8a.1^rk29/rk29^* and *aqp1a.1^rk28/rk28^;aqp8a.1^rk29/rk29^*zebrafish raised in *Tg(fli1:Lifeact-mCherry)^ncv7^;(fli1a:H2B-EGFP)*^ncv69^ transgenic line or in *Tg(fli1ep:aqp1a.1-mEmerald)^rk30^;(fli1:myr-mCherry)^ncv1^*and *Tg(fli1ep:aqp8a.1-mEmerald)^rk31^; (fli1:myr-mCherry)^ncv1^*zebrafish. Confocal z-stacks images of embryos were taken at 50 - 54 hpf and 3 dpf using Olympus UPLSAPO 40x/NA 1.25 silicone oil immersion objective. ISVs no. 5 to 15 were used for quantification. For ISV diameter, 5 measurements were made along the length of each ISV using ImageJ (NIH) and the average was plotted.

### Chemical treatment

Latrunculin B (Merck Millipore) was dissolved in DMSO to 1 mg/ml and stored at -20^0^C. Ki8751 (Selleck Chemicals) was prepared as 5 mM solution in DMSO and stored at -80^0^C. DCPIB (Tocris) was prepared as 10 mM solution in EtOH and stored at -20^0^C. All compounds were diluted to the desired concentration in E3 medium (see in figure legends). Embryos were treated from 20 hpf for 6 (Ki8751 and DCPIB) or 8 hours (Latrunculin B). For imaging experiments, the same concentrations of chemicals were added to the agarose and E3 medium.

### Cell culture

Adult human aortic endothelial cells (HAEC) (Lonza, lot-20TL231227) were cultured in EGM medium (Lonza) and used at passage 3. Cells were seeded in EBM medium (Lonzo) supplemented with 2% FBS (Gibco) at 4 x 10^5^ cells/well on 24-well plate (Trueline) coated with 5 μg/ml fibronectin (Sigma-Aldrich). Cells were treated with either 0.01% DMSO or different concentrations of Ki8751 inhibitor for 6 hours. Cells were lysed with TRI reagent and RNA was isolated using Direct-zol RNA MicroPrep kit (Zymo Research).

### Single-cell RNA sequencing

ECs were isolated from *Tg(kdrl:EGFP)^s843^* transgenic embryos. Cell sorting was carried out with FACSAriaII Cell Sorter (BD Bioscience). Single-cell suspension was loaded into the 10X Chromium system and cDNA libraries were constructed using Chromium Next GEM Single Cell 3′ GEM, Library and Gel Bead Kit v2 (10X Genomics) according to manufacturer’s protocol. Library was sequenced on the Illumina HiSeq 1500 Sequencer (Illumina, USA). Cell Ranger v2.1 was used to de-multiplex raw base call (BCL) files generated by Illumina sequencers into FASTQ files.

### Single-cell RNA sequencing data analysis

The raw sequence data from above mentioned internal sequencing and also a public zebrafish embryo data at 24 hpf stage (Gurung et al., 2022) were processed. The public dataset was obtained from NCBI GEO database (accession number GSE202912). The raw sequencing reads from both datasets were mapped to the zebrafish genome assembly (GRCz11, Ensembl release 112). Further analyses were performed using Seurat package (version 4.3.0) in R software (https://www.r-project.org). The expression matrices were first filtered by keeping genes that are expressed in a minimum of 3 cells and cells that expressed a minimum of 200 genes for downstream analysis. The data were then normalized using NormalizeData function which normalizes the gene expression for each cell by the total expression counts (with a scale factor 10,000). To correct for batch effect between the data from the two-time points, the rpca method in the Seurat package was applied to integrate the data. The top 2000 variable genes identified using the vst method in FindVariableFeatures function were used for principal component analysis in RunPCA function, and the first 30 principal components were used for visualization analysis with Uniform Manifold Approximation and Projection (UMAP) method, and in FindNeighbors function analysis. The cell clustering resolution was set at 0.5 in FindClusters function. The Seurat object was then processed using the ShinyCell package (version: 2.1.0) for gene expression visualizations.

### Statistical analysis

Statistical analysis was performed using Prism software version 10.2.0 (GraphPad). The variance between the mean values of two groups was evaluated using the unpaired Student’s *t*-test. For assessment of more than three groups, we used one-way analysis of variance (ANOVA) test. A *P* value of <0.05 was considered statistically significant. Statistic details can be found in each figure legend.

## Data availability

The scRNA-seq raw data from this study is available at NCBI’s Gene Expression Omnibus database (accession number GSE262232). The data supporting the fundings of this study are available from the corresponding author upon reasonable request.

## Acknowledgements

We thank members of the Phng Lab, Y-C. Wang and S. Thukral for discussions and suggestions; G. Chen, E. Taniguchi, RIKEN BDR Research Aquarium and RIKEN Kobe BioImaging Facilities & Factory for technical assistance; J. Vermot, M. Francois and A. Yap for comments on the manuscript. This work was supported by core funding from RIKEN BDR (to L.K.P.), RIKEN BDR-Otsuka Pharmaceutical Collaboration Center (I.K.), the Naito Foundation (L.K.P.), the JSPS Grants-in-Aid for Scientific Research (KAKENHI) grants (22H02624, 22H05168 to L.K.P; 22K06244 to I.K.), Swedish Research Council (2015-00550 to C.B.); Swedish Cancer Society (2018/449, 2018/1154 to C.B.); Knut and Alice Wallenberg Foundation (2020.0057, 2018.0218 to C.B.); Swedish Brain Foundation (ALZ2019-0130, ALZ2022-0005 to C.B.); the Leducq Foundation (22CVD01, 23CVD02 to C.B.) and the Innovative Medicines Initiative (IM2PACT-807015 to C.B.)

## Authorship contributions

Conceptualization: L.K.P.; Methodology: I.K., L.H., H.W. and L.K.P.; Formal analysis: I.K. and L.H.; Resources: C.B. and L.K.P.; Writing: I.K. and L.K.P; Supervision: L.K.P.

## References

Agre P, King LS, Yasui M, Guggino WB, Ottersen OP, Fujiyoshi Y, Engel A, Nielsen S. 2002. Aquaporin water channels – from atomic structure to clinical medicine. Journal of Physiology 542:3–16. doi:10.1113/jphysiol.2002.020818

Ando K, Wang W, Peng D, Chiba A, Lagendijk AK, Barske L, Crump JG, Stainier DYR, Lendahl U, Koltowska K, Hogan BM, Fukuhara S, Mochizuki N, Betsholtz C. 2019. Peri-arterial specification of vascular mural cells from naïve mesenchyme requires Notch signaling. Development 146:dev165589-13. doi:10.1242/dev.165589

Bill BR, Korzh V. 2014. Choroid plexus in developmental and evolutionary perspective. Frontiers in Neuroscience 8:363. doi:10.3389/fnins.2014.00363

Boer LL de, Vanes L, Melgrati S, O’May JB, Hayward D, Driscoll PC, Day J, Griffiths A, Magueta R, Morrell A, MacRae JI, Köchl R, Tybulewicz VLJ. 2023. T cell migration requires ion and water influx to regulate actin polymerization. Nature Communications 14:7844. doi:10.1038/s41467-023-43423-8

Calamita G, Ferri D, Gena P, Liquori GE, Cavalier A, Thomas D, Svelto M. 2005. The Inner Mitochondrial Membrane Has Aquaporin-8 Water Channels and Is Highly Permeable to Water*. Journal of Biological Chemistry 280:17149–17153. doi:10.1074/jbc.c400595200

Chae YK, Woo J, Kim M-J, Kang SK, Kim MS, Lee J, Lee SK, Gong G, Kim YH, Soria JC, Jang SJ, Sidransky D, Moon C. 2008. Expression of Aquaporin 5 (AQP5) Promotes Tumor Invasion in Human Non Small Cell Lung Cancer. PLoS ONE 3:e2162. doi:10.1371/journal.pone.0002162

Chan CJ, Costanzo M, Ruiz-Herrero T, Mönke G, Petrie RJ, Bergert M, Diz-Muñoz A, Mahadevan L, Hiiragi T. 2019. Hydraulic control of mammalian embryo size and cell fate. Nature 571:112–116. doi:10.1038/s41586-019-1309-x

Chen C, Qin Y, Xu Y, Wang X, Lei W, Shen X, Chen L, Wang L, Gong J, Wang Y, Hu S, Liu D. 2024. Aquaporins enriched in endothelial vacuole membrane regulate the diameters of microvasculature in hyperglycaemia. Cardiovascular Research 120:1065–1080. doi:10.1093/cvr/cvae085

Choudhury MI, Benson MA, Sun SX. 2022. Trans-epithelial fluid flow and mechanics of epithelial morphogenesis. Seminars in Cell & Developmental Biology 131:146–159. doi:10.1016/j.semcdb.2022.05.020

Chugh M, Munjal A, Megason SG. 2022. Hydrostatic pressure as a driver of cell and tissue morphogenesis. Seminars in Cell & Developmental Biology 131:134–145. doi:10.1016/j.semcdb.2022.04.021

Clarke DN, Martin AC. 2021. Actin-based force generation and cell adhesion in tissue morphogenesis. Current Biology 31:R667–R680. doi:10.1016/j.cub.2021.03.031

Conner MT, Conner AC, Bland CE, Taylor LHJ, Brown JEP, Parri HR, Bill RM. 2012. Rapid Aquaporin Translocation Regulates Cellular Water Flow. Journal of Biological Chemistry 287:11516–11525. doi:10.1074/jbc.m111.329219

Day RE, Kitchen P, Owen DS, Bland C, Marshall L, Conner AC, Bill RM, Conner MT. 2014. Human aquaporins: regulators of transcellular water flow. Biochimica et Biophysica Acta 1840:1492–1506. doi:10.1016/j.bbagen.2013.09.033

Dumortier JG, Verge-Serandour ML, Tortorelli AF, Mielke A, Plater L de, Turlier H, Maître J-L. 2019. Hydraulic fracturing and active coarsening position the lumen of the mouse blastocyst. Science 365:465–468. doi:10.1126/science.aaw7709

Emelyanov A, Gao Y, Naqvi NI, Parinov S. 2006. Trans-Kingdom Transposition of the Maize Dissociation Element. Genetics 174:1095–1104. doi:10.1534/genetics.106.061184

Farinas J, Kneen M, Moore M, Verkman AS. 1997. Plasma Membrane Water Permeability of Cultured Cells and Epithelia Measured by Light Microscopy with Spatial Filtering. Journal of General Physiology 110:283–296. doi:10.1085/jgp.110.3.283

Figueiredo AM, Barbacena P, Russo A, Vaccaro S, Ramalho D, Pena A, Lima AP, Ferreira RR, Fidalgo MA, El-Marjou F, Carvalho Y, Vasconcelos FF, Lennon-Duménil A-M, Vignjevic DM, Franco CA. 2021. Endothelial cell invasion is controlled by dactylopodia. Proceedings of the National Academy of Sciences 118:e2023829118. doi:10.1073/pnas.2023829118

Fukuhara S, Zhang J, Yuge S, Ando K, Wakayama Y, Sakaue-Sawano A, Miyawaki A, Mochizuki N. 2014. Visualizing the cell-cycle progression of endothelial cells in zebrafish. Developmental Biology 393:10–23. doi:10.1016/j.ydbio.2014.06.015

Gagnon JA, Valen E, Thyme SB, Huang P, Akhmetova L, Ahkmetova L, Pauli A, Montague TG, Zimmerman S, Richter C, Schier AF. 2014. Efficient mutagenesis by Cas9 protein-mediated oligonucleotide insertion and large-scale assessment of single-guide RNAs. PLoS ONE 9:e98186. doi:10.1371/journal.pone.0098186

Gebala V, Collins R, Geudens I, Phng L-K, Gerhardt H. 2016. Blood flow drives lumen formation by inverse membrane blebbing during angiogenesis in vivo. Nature Cell Biology 18:443–450. doi:10.1038/ncb3320

Gerhardt H, Golding M, Fruttiger M, Ruhrberg C, Lundkvist A, Abramsson A, Jeltsch M, Mitchell C, Alitalo K, Shima D, Betsholtz C. 2003. VEGF guides angiogenic sprouting utilizing endothelial tip cell filopodia. Journal of Cell Biology 161:1163–1177. doi:10.1083/jcb.200302047

Gradilone SA, Carreras FI, Lehmann GL, Marinelli RA. 2005. Phosphoinositide 3-kinase is involved in the glucagon-induced translocation of aquaporin-8 to hepatocyte plasma membrane. Biology of the Cell 97:831–836. doi:10.1042/bc20040115

Gradilone SA, García F, Huebert RC, Tietz PS, Larocca MC, Kierbel A, Carreras FI, Larusso NF, Marinelli RA. 2003. Glucagon induces the plasma membrane insertion of functional aquaporin-8 water channels in isolated rat hepatocytes. Hepatology 37:1435–1441. doi:10.1053/jhep.2003.50241

Gross-Thebing T, Paksa A, Raz E. 2014. Simultaneous high-resolution detection of multiple transcripts combined with localization of proteins in whole-mount embryos. BMC Biology 12:55. doi:10.1186/s12915-014-0055-7

Gunasekar SK, Xie L, Kumar A, Hong J, Chheda PR, Kang C, Kern DM, My-Ta C, Maurer J, Heebink J, Gerber EE, Grzesik WJ, Elliot-Hudson M, Zhang Y, Key P, Kulkarni CA, Beals JW, Smith GI, Samuel I, Smith JK, Nau P, Imai Y, Sheldon RD, Taylor EB, Lerner DJ, Norris AW, Klein S, Brohawn SG, Kerns R, Sah R. 2022. Small molecule SWELL1 complex induction improves glycemic control and nonalcoholic fatty liver disease in murine Type 2 diabetes. Nature Communications 13:784. doi:10.1038/s41467-022-28435-0

Gurung S, Restrepo NK, Chestnut B, Klimkaite L, Sumanas S. 2022. Single-cell transcriptomic analysis of vascular endothelial cells in zebrafish embryos. Scientific Reports 12:13065. doi:10.1038/s41598-022-17127-w

Herrera M, Garvin JL. 2007. Novel role of AQP-1 in NO-dependent vasorelaxation. American Journal of Physiology-Renal Physiology 292:F1443–F1451. doi:10.1152/ajprenal.00353.2006

Herrera M, Hong NJ, Garvin JL. 2006. Aquaporin-1 Transports NO Across Cell Membranes. Hypertension 48:157–164. doi:10.1161/01.hyp.0000223652.29338.77

Herwig L, Blum Y, Krudewig A, Ellertsdottir E, Lenard A, Belting H-G, Affolter M. 2011. Distinct cellular mechanisms of blood vessel fusion in the zebrafish embryo. Current Biology 21:1942–1948. doi:10.1016/j.cub.2011.10.016

Hogan BM, Bos FL, Bussmann J, Witte M, Chi NC, Duckers HJ, Schulte-Merker S. 2009. ccbe1 is required for embryonic lymphangiogenesis and venous sprouting. Nature Genetics 41:396–398. doi:10.1038/ng.321

Jahn TP, Møller ALB, Zeuthen T, Holm LM, Klærke DA, Mohsin B, Kühlbrandt W, Schjoerring JK. 2004. Aquaporin homologues in plants and mammals transport ammonia. FEBS Letters 574:31–36. doi:10.1016/j.febslet.2004.08.004

Jin S-W, Beis D, Mitchell T, Chen J-N, Stainier DY. 2005. Cellular and molecular analyses of vascular tube and lumen formation in zebrafish. Development 132: 5199–5209. doi:10.1242/dev.02087

Jung HJ, Park J-Y, Jeon H-S, Kwon T-H. 2011. Aquaporin-5: A Marker Protein for Proliferation and Migration of Human Breast Cancer Cells. PLoS ONE 6:e28492. doi:10.1371/journal.pone.0028492

Karlsson T, Bolshakova A, Magalhães MAO, Loitto VM, Magnusson K-E. 2013. Fluxes of Water through Aquaporin 9 Weaken Membrane-Cytoskeleton Anchorage and Promote Formation of Membrane Protrusions. PLoS ONE 8:e59901. doi:10.1371/journal.pone.0059901

Kimmel CB, Ballard WW, Kimmel SR, Ullmann B, Schilling TF. 1995. Stages of embryonic development of the zebrafish. Developmental dynamics 203:253--310. doi:10.1002/aja.1002030302

Kondrychyn I, Garcia-Lecea M, Emelyanov A, Parinov S, Korzh V. 2009. Genome-wide analysis of Tol2 transposon reintegration in zebrafish. BMC Genomics 10:418–418. doi:10.1186/1471-2164-10-418

Kondrychyn I, Kelly DJ, Carretero NT, Nomori A, Kato K, Chong J, Nakajima H, Okuda S, Mochizuki N, Phng L-K. 2020. Marcksl1 modulates endothelial cell mechanoresponse to haemodynamic forces to control blood vessel shape and size. Nature Communications 11:5476–18. doi:10.1038/s41467-020-19308-5

Koun S, Kim J-D, Rhee M, Kim M-J, Huh T-L. 2016. Spatiotemporal expression pattern of the zebrafish aquaporin 8 family during early developmental stages. Gene Expression Patterns 21:1–6. doi:10.1016/j.gep.2016.06.001

Kozono D, Yasui M, King LS, Agre P. 2002. Aquaporin water channels: atomic structure molecular dynamics meet clinical medicine. Journal of Clinical Investigation 109:1395–1399. doi:10.1172/jci15851

Kwon H-B, Wang S, Helker CSM, Rasouli SJ, Maischein H-M, Offermanns S, Herzog W, Stainier DYR. 2016. In vivo modulation of endothelial polarization by Apelin receptor signalling. Nature Communications 7:11805–12. doi:10.1038/ncomms11805

Li Y, Konstantopoulos K, Zhao R, Mori Y, Sun SX. 2020. The importance of water and hydraulic pressure in cell dynamics. Journal of Cell Science 133:jcs240341. doi:10.1242/jcs.240341

Liu K, Nagase H, Huang CG, Calamita G, Agre P. 2006. Purification and functional characterization of aquaporin-8. Biology of the Cell 98:153–161. doi:10.1042/bc20050026

Loitto VM, Huang C, Sigal YJ, Jacobson K. 2007. Filopodia are induced by aquaporin-9 expression. Experimental Cell Research 313:1295–1306. doi:10.1016/j.yexcr.2007.01.023

Loitto VM, Karlsson T, Magnusson K. 2009. Water flux in cell motility: Expanding the mechanisms of membrane protrusion. Cell Motility and the Cytoskeleton 66:237–247. doi:10.1002/cm.20357

Ma T, Yang B, Verkman AS. 1997. Cloning of a Novel Water and Urea-Permeable Aquaporin from Mouse Expressed Strongly in Colon, Placenta, Liver, and Heart. Biochemical and Biophysical Research Communications 240:324–328. doi:10.1006/bbrc.1997.7664

Maltaneri RE, Schiappacasse A, Chamorro ME, Nesse AB, Vittori DC. 2020. Aquaporin-1 plays a key role in erythropoietin-induced endothelial cell migration. Biochimica et Biophysica Acta (BBA) - Molecular Cell Research 1867:118569. doi:10.1016/j.bbamcr.2019.118569

Markou A, Unger L, Abir-Awan M, Saadallah A, Halsey A, Balklava Z, Conner M, Törnroth-Horsefield S, Greenhill SD, Conner A, Bill RM, Salman MM, Kitchen P. 2022. Molecular mechanisms governing aquaporin relocalisation. Biochimica et Biophysica Acta (BBA) - Biomembranes 1864:183853. doi:10.1016/j.bbamem.2021.183853

McLennan R, McKinney MC, Teddy JM, Morrison JA, Kasemeier-Kulesa JC, Ridenour DA, Manthe CA, Giniunaite R, Robinson M, Baker RE, Maini PK, Kulesa PM. 2019. Neural crest cells bulldoze through the microenvironment using Aquaporin 1 to stabilize filopodia. Development 147:dev185231. doi:10.1242/dev.185231

Meeker ND, Hutchinson SA, Ho L, Trede NS. 2007. Method for isolation of PCR-ready genomic DNA from zebrafish tissues. Biotechniques 43:610–614. doi:10.2144/000112619

Mosaliganti KR, Swinburne IA, Chan CU, Obholzer ND, Green AA, Tanksale S, Mahadevan L, Megason SG. 2019. Size control of the inner ear via hydraulic feedback. eLife 8:e39596. doi:10.7554/elife.39596

Munjal A, Lecuit T. 2014. Actomyosin networks and tissue morphogenesis. Development 141:1789–1793. doi: 10.1242/dev.091645

Murrell M, Oakes PW, Lenz M, Gardel ML. 2015. Forcing cells into shape: the mechanics of actomyosin contractility. Nature Cell Biology 16:486–498. doi:10.1038/nrm4012

Nagy TL, Strickland J, Weiner OD. 2023. Neutrophils actively swell to potentiate rapid migration. bioRxiv 2023.05.15.540704. doi:10.1101/2023.05.15.540704

Nakajima H, Yamamoto K, Agarwala S, Terai K, Fukui H, Fukuhara S, Ando K, Miyazaki T, Yokota Y, Schmelzer E, Belting H-G, Affolter M, Lecaudey V, Mochizuki N. 2017. Flow-Dependent Endothelial YAP Regulation Contributes to Vessel Maintenance. Developmental Cell 40:523–536.e6. doi:10.1016/j.devcel.2017.02.019

Nakhoul NL, Davis BA, Romero MF, Boron WF. 1998. Effect of expressing the water channel aquaporin-1 on the CO2 permeability of Xenopus oocytes. American Journal of Physiology-Cell Physiology 274:C543–C548. doi:10.1152/ajpcell.1998.274.2.c543

Parab S, Quick RE, Matsuoka RL. 2021. Endothelial cell-type-specific molecular requirements for angiogenesis drive fenestrated vessel development in the brain. eLife 10:e64295. doi:10.7554/elife.64295

Pfaffl MW. 2001. A new mathematical model for relative quantification in real-time RT–PCR. Nucleic Acids Research 29:e45–e45. doi:10.1093/nar/29.9.e45

Phng L-K, Stanchi F, Gerhardt H. 2013. Filopodia are dispensable for endothelial tip cell guidance. Development 140:4031–4040. doi:10.1242/dev.097352

Prasad GV, Coury LA, Finn F, Zeidel ML. 1998. Reconstituted aquaporin 1 water channels transport CO2 across membranes. Journal of Biological Chemistry 273:33123–33126. doi:10.1074/jbc.273.50.33123

Preston GM, Carroll TP, Guggino WB, Agre P. 1992. Appearance of water channels in Xenopus oocytes expressing red cell CHIP28 protein. Science 256:385–387. doi:10.1126/science.256.5055.385

Qiu Z, Dubin AE, Mathur J, Tu B, Reddy K, Miraglia LJ, Reinhardt J, Orth AP, Patapoutian A. 2014. SWELL1, a Plasma Membrane Protein, Is an Essential Component of Volume-Regulated Anion Channel. Cell 157:447–458. doi:10.1016/j.cell.2014.03.024

Rehn K, Wong KS, Balciunas D, Sumanas S. 2011. Zebrafish enhancer trap line recapitulates embryonic aquaporin 1a expression pattern in vascular endothelial cells. International Journal of Developmental Biology 55:613–618. doi:10.1387/ijdb.103249kp

Saadoun S, Papadopoulos MC, Hara-Chikuma M, Verkman AS. 2005. Impairment of angiogenesis and cell migration by targeted aquaporin-1 gene disruption. Nature 434:786– 792. doi:10.1038/nature03460

Sauteur L, Affolter M, Belting H-G. 2017. Distinct and redundant functions of Esama and VE-cadherin during vascular morphogenesis. Development 144:1554–1565. doi:10.1242/dev.140038

Siekmann AF, Affolter M, Belting H-G. 2013. The tip cell concept 10 years after: New players tune in for a common theme. Experimental Cell Research 319:1255–1263. doi:10.1016/j.yexcr.2013.01.019

Soria LR, Fanelli E, Altamura N, Svelto M, Marinelli RA, Calamita G. 2010. Aquaporin-8-facilitated mitochondrial ammonia transport. Biochemical and Biophysical Research Communications 393:217–221. doi:10.1016/j.bbrc.2010.01.104

Stroka KM, Jiang H, Chen S-H, Tong Z, Wirtz D, Sun SX, Konstantopoulos K. 2014. Water permeation drives tumor cell migration in confined microenvironments. Cell 157:611–623. doi:10.1016/j.cell.2014.02.052

Talbot K, Kwong RWM, Gilmour KM, Perry SF. 2015. The water channel aquaporin-1a1 facilitates movement of CO2 and ammonia in zebrafish (Danio rerio) larvae. Journal of Experimental Biology 218:3931–3940. doi:10.1242/jeb.129759

Thisse C, Thisse B. 2007. High-resolution in situ hybridization to whole-mount zebrafish embryos. Nature Protocols 3:59–69. doi:10.1038/nprot.2007.514

Tingaud-Sequeira A, Calusinska M, Finn RN, Chauvigné F, Lozano J, Cerdà J. 2010. The zebrafish genome encodes the largest vertebrate repertoire of functional aquaporins with dual paralogy and substrate specificities similar to mammals. BMC Evolutionary Biology 10:38. doi:10.1186/1471-2148-10-38

Voss FK, Ullrich F, Münch J, Lazarow K, Lutter D, Mah N, Andrade-Navarro MA, Kries JP von, Stauber T, Jentsch TJ. 2014. Identification of LRRC8 Heteromers as an Essential Component of the Volume-Regulated Anion Channel VRAC. Science 344:634–638. doi:10.1126/science.1252826

Wakayama Y, Fukuhara S, Ando K, Matsuda M, Mochizuki N. 2015. Cdc42 Mediates Bmp-Induced Sprouting Angiogenesis through Fmnl3-Driven Assembly of Endothelial Filopodia in Zebrafish. Developmental Cell 32:109–122. doi:10.1016/j.devcel.2014.11.024

Wang Y, Cohen J, Boron WF, Schulten K, Tajkhorshid E. 2007. Exploring gas permeability of cellular membranes and membrane channels with molecular dynamics. Journal of Structural Biology 157:534–544. doi:10.1016/j.jsb.2006.11.008

Wu DQ, Yang ZF, Wang KJ, Feng XY, Lv ZJ, Li Y, Jian ZX. 2017. AQP8 inhibits colorectal cancer growth and metastasis by down-regulating PI3K/AKT signaling and PCDH7 expression. American Journal of Cancer Research 8:266–279.

Zeidel ML, Ambudkar SV, Smith BL, Agre P. 1992. Reconstitution of functional water channels in liposomes containing purified red cell CHIP28 protein. Biochemistry 31:7436–7440. doi:10.1021/bi00148a002

Zhang Y, Li Y, Thompson KN, Stoletov K, Yuan Q, Bera K, Lee SJ, Zhao R, Kiepas A, Wang Y, Mistriotis P, Serra SA, Lewis JD, Valverde MA, Martin SS, Sun SX, Konstantopoulos K. 2022. Polarized NHE1 and SWELL1 regulate migration direction, efficiency and metastasis. Nature Communications 13:6128. doi:10.1038/s41467-022-33683-1

