## Supplemental figures, table and movie legends for "Combined forces of hydrostatic pressure and actin polymerization drive endothelial tip cell migration and sprouting angiogenesis"

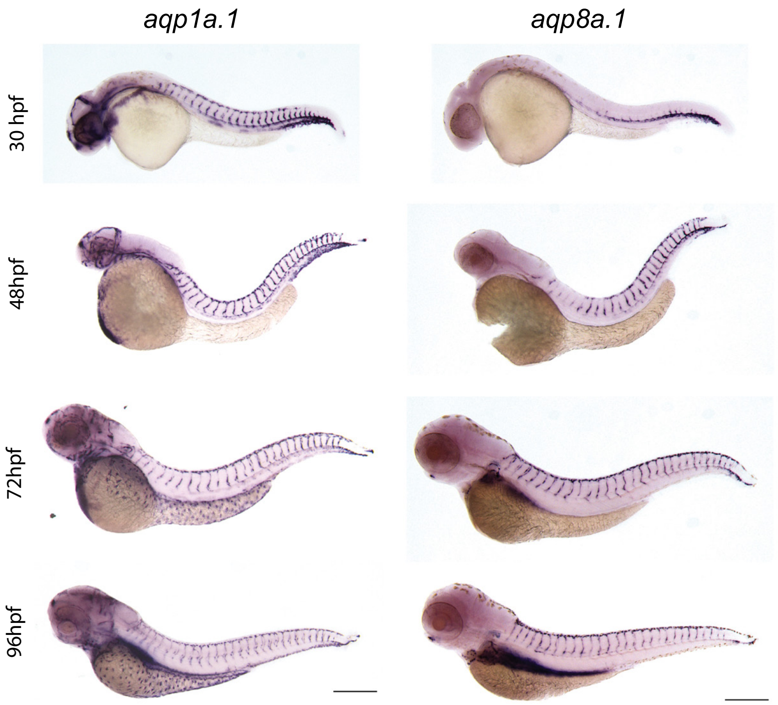

**Figure S1. Expression of endothelial-specific *aquaporins* during development.**
Whole mount *in situ* hybridization with *aqp1a.1* and *aqp8a.1* RNA probes at different developmental stages. Images are representative of 10 embryos for every stage and probe (n=2 independent experiments). Scale bar, 250 µm.

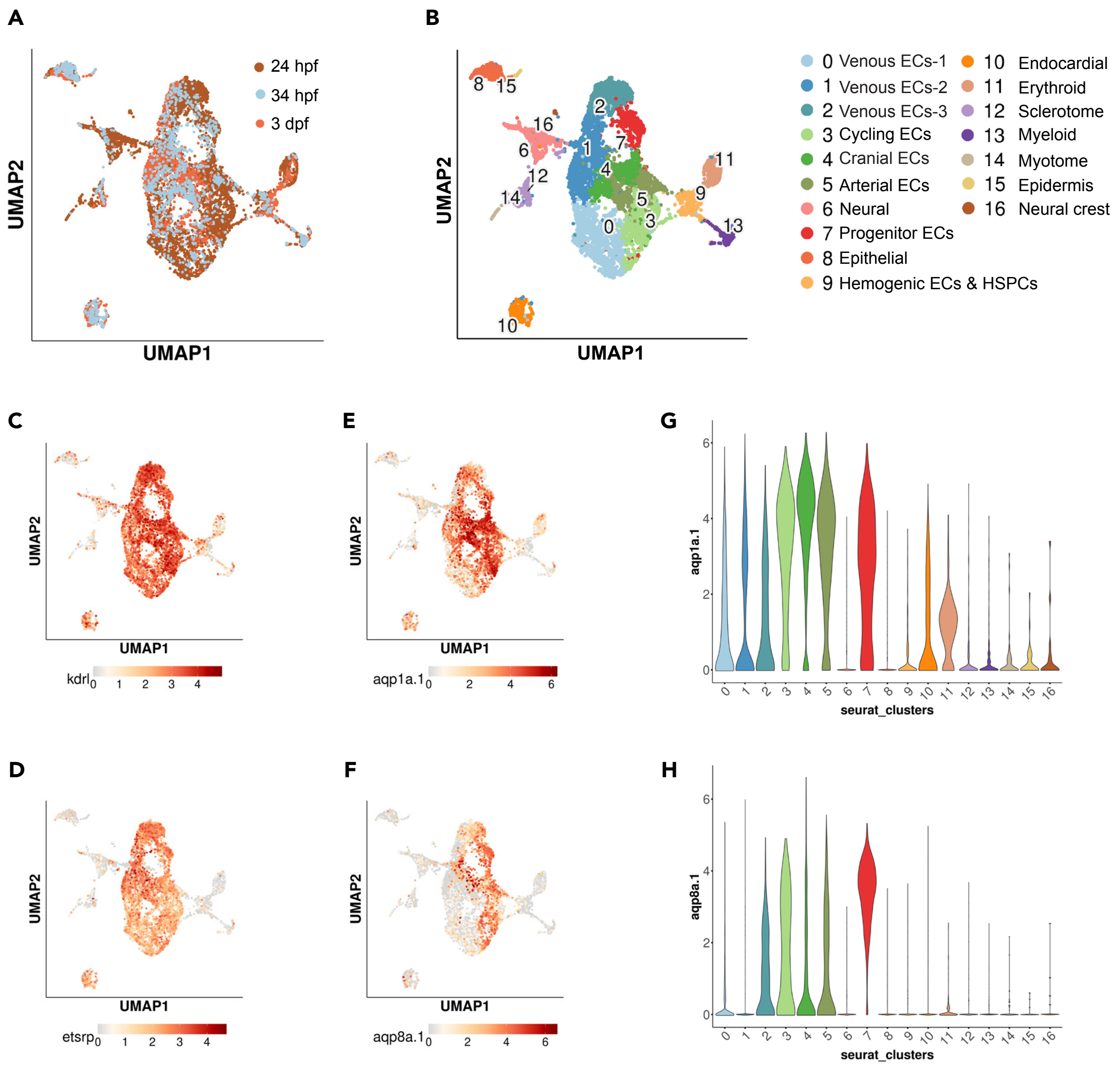

**Figure S2. scRNA-seq analysis of *kdrl* positive cells from 24 hpf, 34 hpf and 3 dpf zebrafish.**
(**A-B**) UMAP plots of 5576 cells from 24 hpf *Tg(kdrl:mCherry)* (Gurung et al., 2022), 1475 cells from 34 hpf *Tg(kdrl:EGFP)* zebrafish and 1692 cells from 3 dpf *Tg(kdrl:EGFP)* zebrafish (A) showing 16 distinct clusters (B). Classifications were based on previously known marker genes that are enriched in each cluster. (**C-F**) UMAP feature plots showing expression of *kdrl* (C), *etsrp*/*etv2* (D), *aqp1a.1* (E) and *aqp8a.1* (F). (**G-H**) Violin plots showing expression of *aqp1a.1* (G) and *aqp8a.1* (H) in different cell populations.

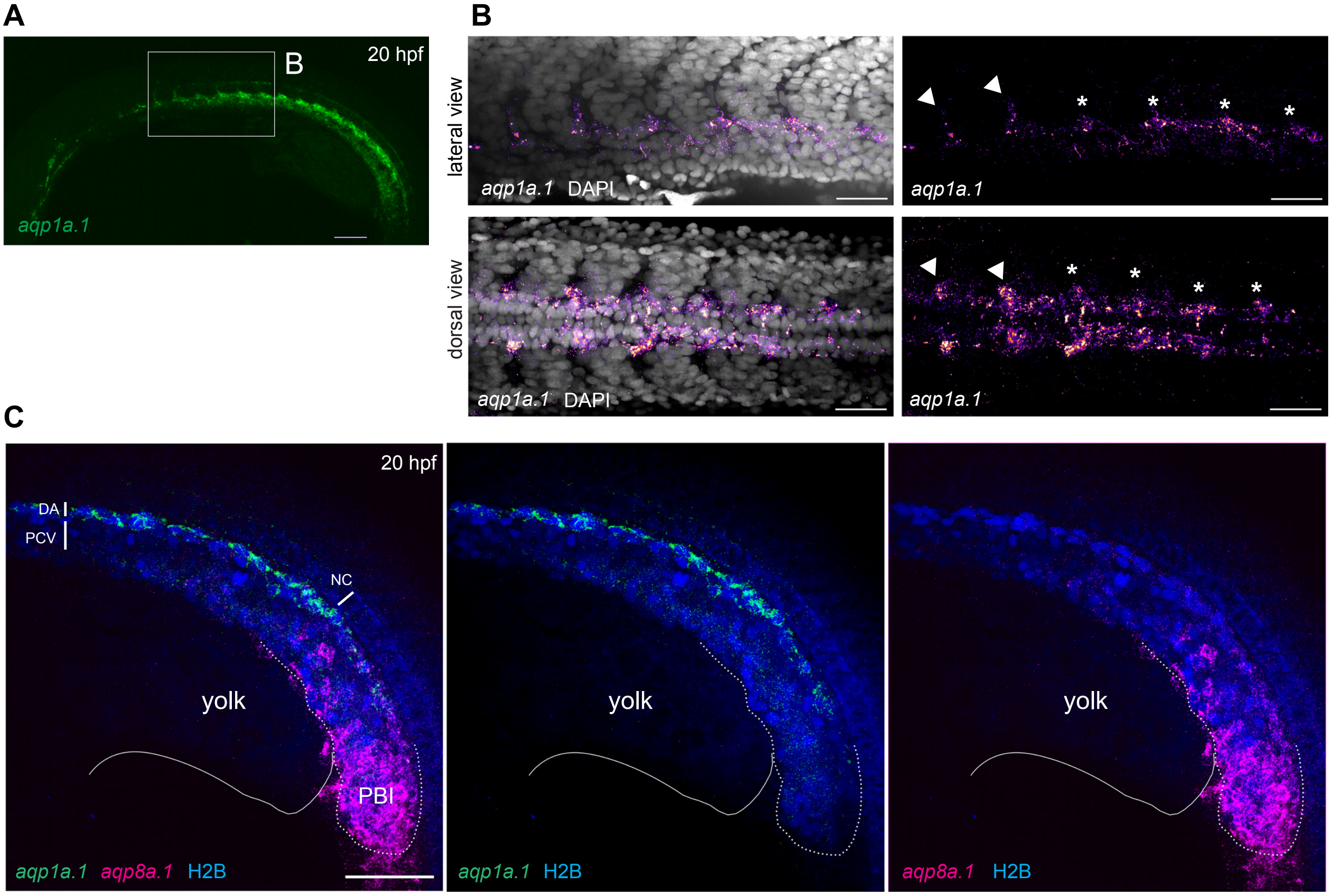

**Figure S3. *aqp1a.1* and *aqp8a.1* are differentially expressed in the dorsal aorta at 20 hpf.**
(**A-B**) *aqp1a.1* mRNA expression in the DA, as determined by RNAscope *in situ* hybridization. B, a magnified region of the anterior trunk. *aqp1a.1* expression is heterogeneous, with higher expression level in the sprouting cells (arrowhead) and in endothelial tip cells that will protrude from the DA (asterisk). (**C**) *aqp8a.1* mRNA expression in the posterior blood island, as determined by RNAscope *in situ* hybridization. Endothelial nuclei are labelled with EGFP (pseudocolored in blue) in *Tg(fli1a:H2B-EGFP)*$^{nvc69}$ transgenic line. Representative confocal images of 9 embryos (n=2 independent experiments). Anterior to the left. DA, dorsal aorta; NC, notochord; PCV, posterior cardinal vein; PBI, posterior blood island. Scale bar, 100 µm (A) and 40 µm (B and C).

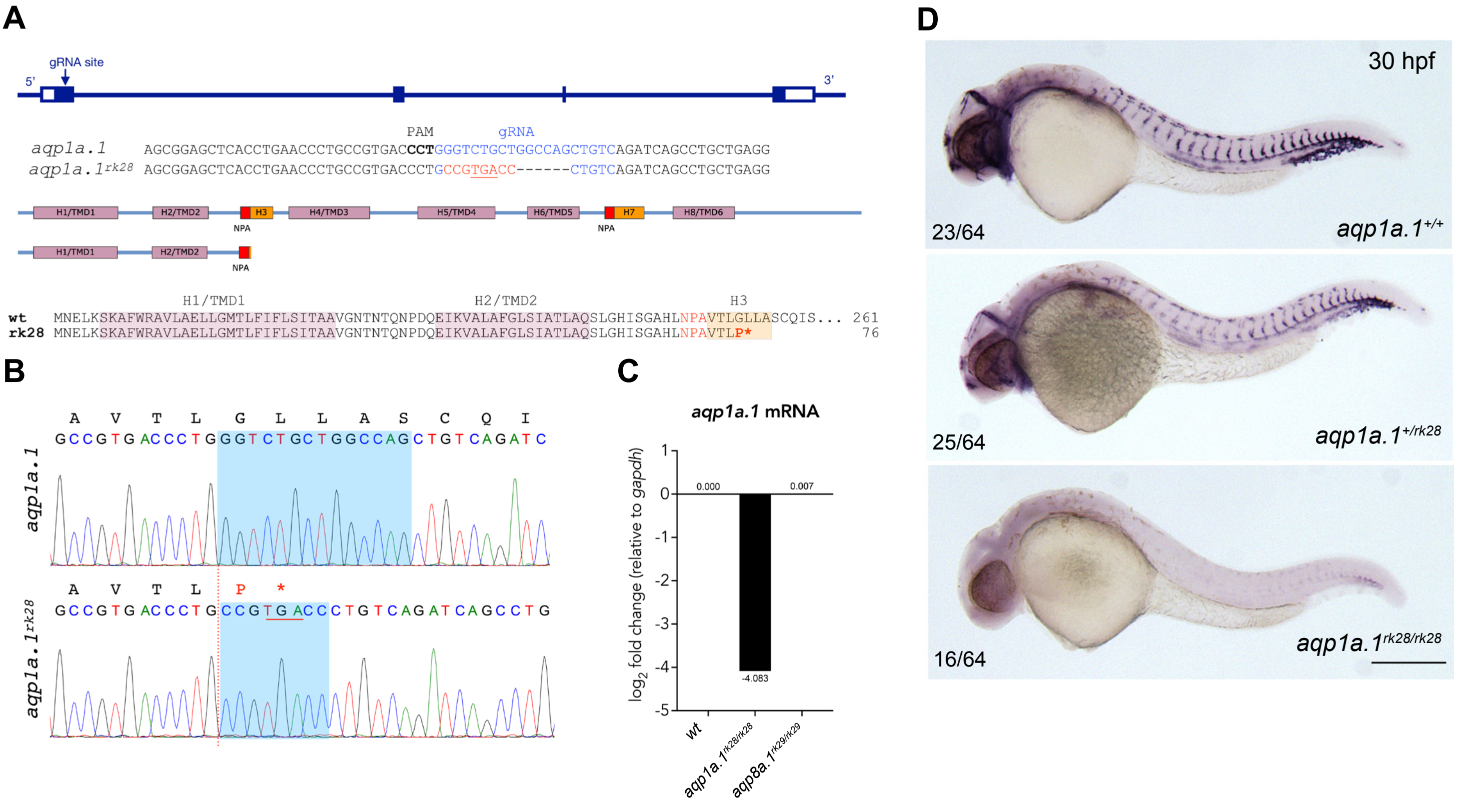

**Figure S4. CRISPR/Cas9-induced mutation in zebrafish *aqp1a.1* gene.**
(**A**) Zebrafish *aqp1a.1* gene structure, gRNA binding site (in blue), *aqp1a.1*$^{rk28}$ allele, Aqp1a.1 wild type (261 aa) and Aqp1a.1$^{rk28}$ (truncated at 76 aa) protein structure. The *rk28* mutation causes a 14-nt deletion and an 8-nt insertion (in red) which leads to a premature termination codon (underlined), and as a result to the loss of 4 out of 6 transmembrane domains/helices (H4/TMD3-H8/TMD6) and 2 membrane-inserted non-membrane-spanning helices (H3 and H7). (**B**) Sequence read showing 14-nt deletion (shadowed area in wild type allele) and 8-nt insertion (shadowed area in *rk28* allele) in *aqp1a.1*$^{rk28}$ allele. Premature termination codon is underlined. Amino acid sequence is shown above the nucleotide sequences. (**C**) qPCR analysis of *aqp1a.1* mRNA expression in wild type, *aqp1a.1*$^{rk28/rk28}$ and *aqp8a.1*$^{rk29/rk29}$ embryos. (**D**) Representative images of wild type, heterozygote and homozygote embryos after whole mount *in situ* hybridization with *aqp1a.1* RNA probe at 30 hpf. Scale bar, 250 μm.

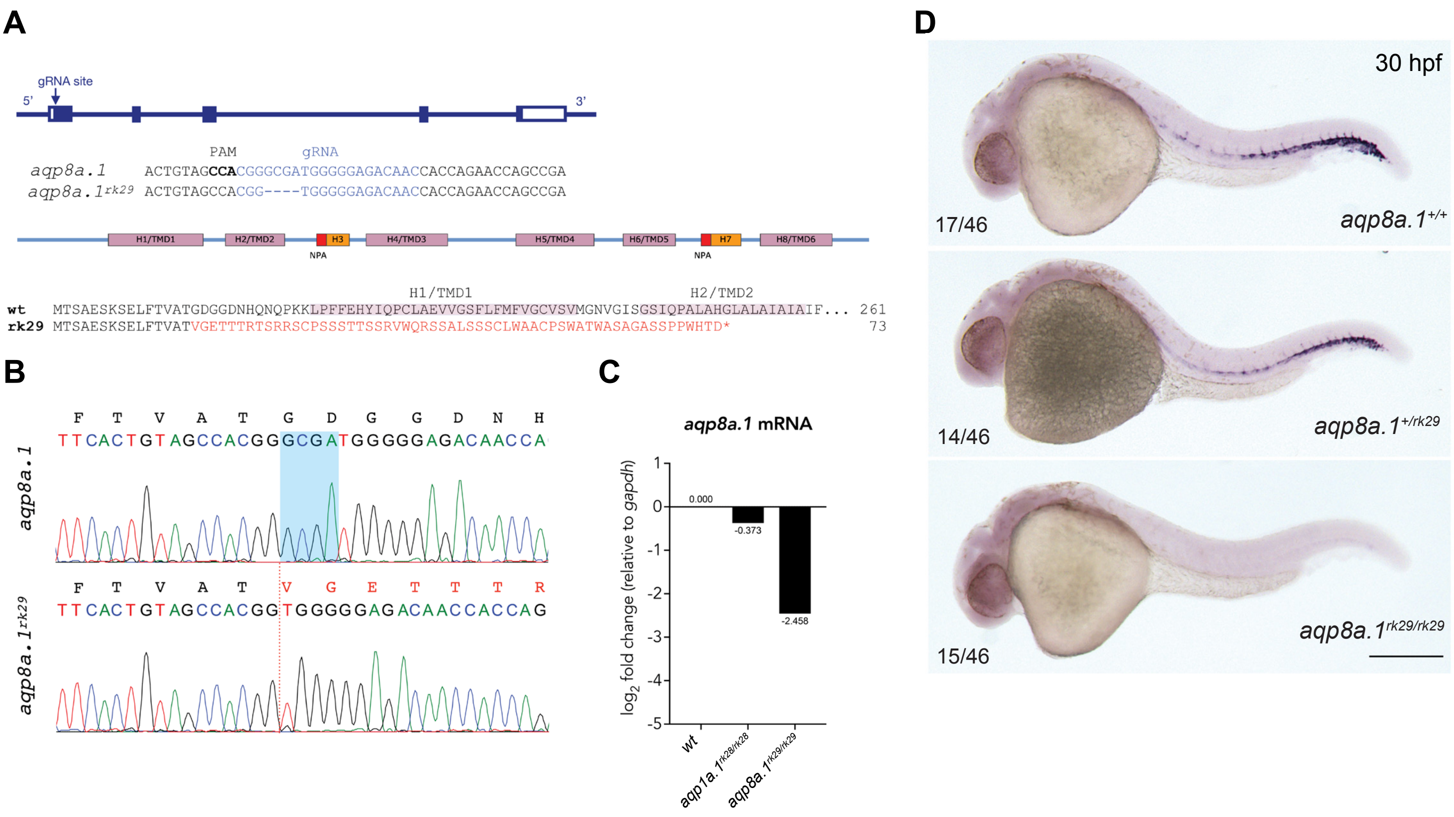

**Figure S5. CRISPR/Cas9-induced mutation in zebrafish *aqp8a.1* gene.**

(**A**) Zebrafish *aqp8a.1* gene structure, gRNA binding site (in blue), *aqp8a.1*$^{rk29}$ allele, Aqp8a.1 wild type (261 aa) and Aqp8a.1$^{rk29}$ (truncated at 73 aa) protein structure. The *rk29* mutation causes a 4-nt deletion which leads to a frameshift after T15 and premature termination codon at amino acid 73 after 58 missense amino acids, and as a result to the loss of 6 transmembrane domains/helices (H1/TMD1-H8/TMD6) and 2 membrane-inserted non-membrane-spanning helices (H3 and H7). (**B**) Sequence read showing 4-nt deletion (shadowed area in wild type allele) in *aqp8a.1*$^{rk29}$ allele (red dotted line). Amino acid sequence is shown above the nucleotide sequences. (**C**) qPCR analysis of *aqp8a.1* mRNA expression in wild type, *aqp1a.1*$^{rk28/rk28}$ and *aqp8a.1*$^{rk29/rk29}$ embryos. (**D**) Representative images of wild type, heterozygote and homozygote embryos after whole mount *in situ* hybridization with *aqp8a.1* RNA probe at 30 hpf. Scale bar, 250 μm.

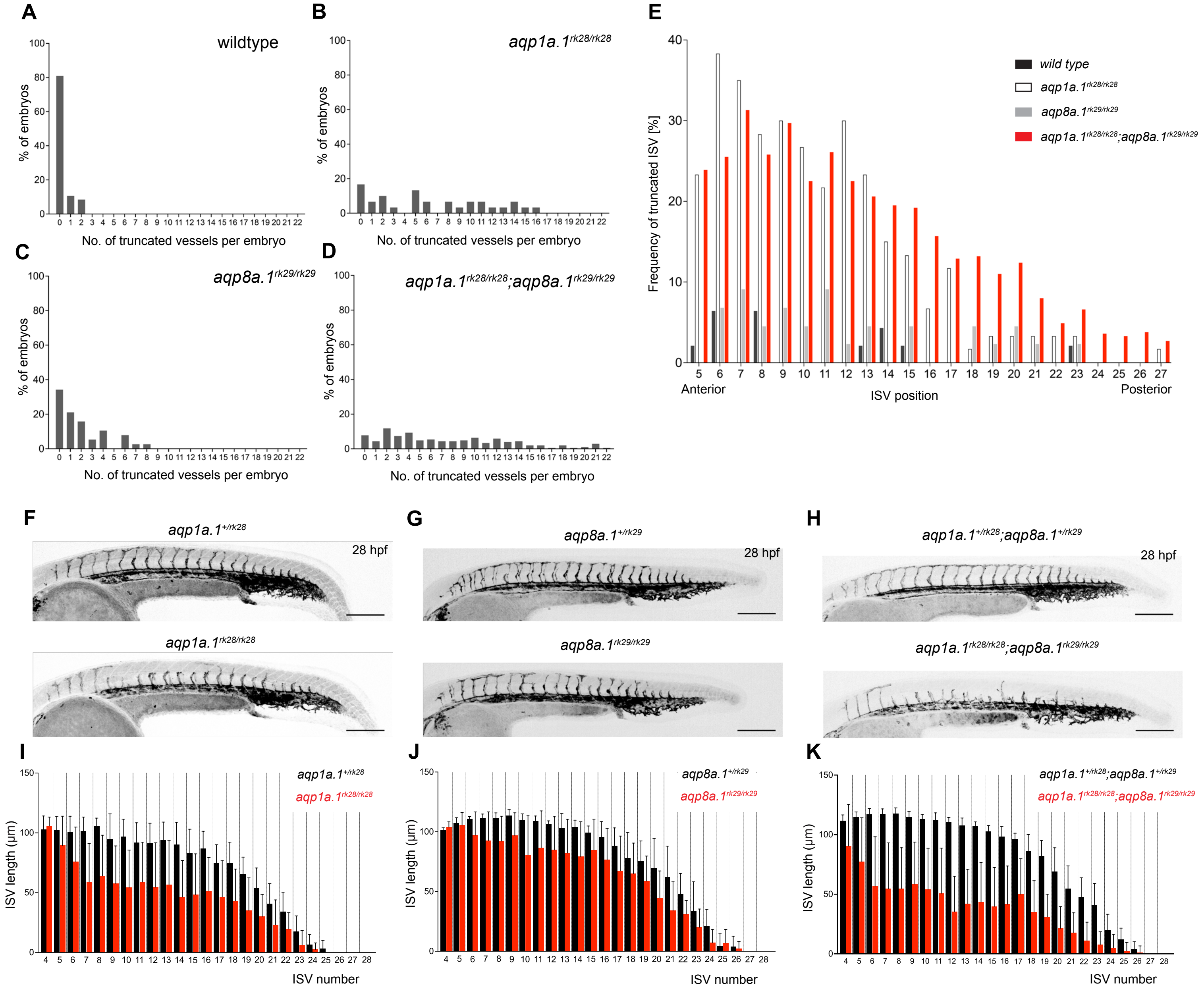

**Figure S6. Phenotype of *aquaporin* mutant embryos.**
(**A-D**) Number of truncated ISVs per embryo in wild type (A, n=47), *aqp1a.1$^{rk28/rk28}$* (B, n=30), *aqp8a.1$^{rk29/rk29}$* (C, n=38) and *aqp1a.1$^{rk28/rk28}$;aqp8a.1$^{rk29/rk29}$* (D, n=204) larvae at 3 dpf. (**E**) Anterior-to-posterior distribution of truncated ISVs in the trunk and tail region. (**F-H**) Maximum intensity projection confocal z-stacks of *aquaporin* heterozygote and homozygote embryos at 28 hpf. Images are representative of 10 embryos for each genotype. Scale bar, 200 µm. (**I-K**) Quantification of ISV length at each position along the length of the embryos in *aquaporin* heterozygote and homozygote embryos at 28 hpf. ISVs, intersegmental vessels.

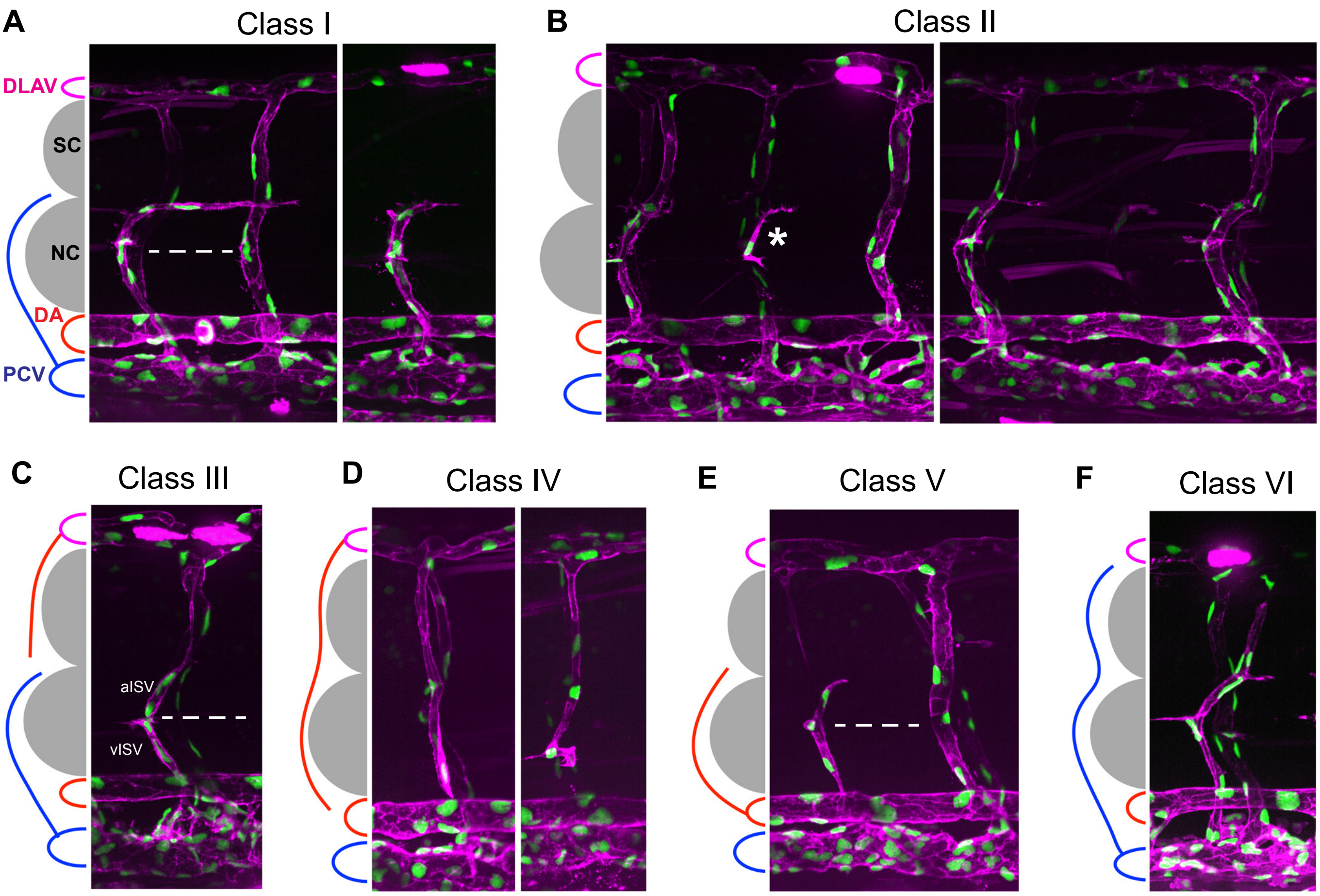

**Figure S7. Vessel phenotype classification in *aqp1a.1*$^{rk28/rk28}$*;aqp8a.1*$^{rk29/rk29}$ mutant embryos at 3 dpf.**

(**A**) Class I (75.2% of truncated ISVs). vISV crosses horizontal myoseptum and turns right (posteriorly towards the tail) along the ventral side of the spinal cord. Vessel comprises 4 ECs.

(**B**) Class II (15.8% of truncated ISVs). Absence of ISV in segmental position (sometimes a single EC is present, which is neither connected to the DA nor DLAV, asterisk).

(**C**) Class III (1.2% of truncated ISVs). vISV does not cross horizontal myoseptum and aISV is not connected to the DA.

(**D**) Class IV (2.2% of truncated ISVs). aISV is not connected to the DA.

(**E**) Class V (2.9% of truncated ISVs). aISV crosses horizontal myoseptum, but has no connection to DLAV. Vessel comprises 3 ECs.

(**F**) Class VI (2.7% of incomplete ISVs). vISV extends over the notochord and spinal cord, but does not connect to DLAV.

In panels A, C and E, dashed line marks an approximate position of horizontal myoseptum. DA, dorsal aorta; DLAV, dorsal longitudinal anastomotic vessel; EC, endothelial cell; ISV, intersegmental vessel; aISV, arterial ISV; vISV, vein ISV; NC, notochord; PCV, posterior cardinal vein; SC, spinal cord.

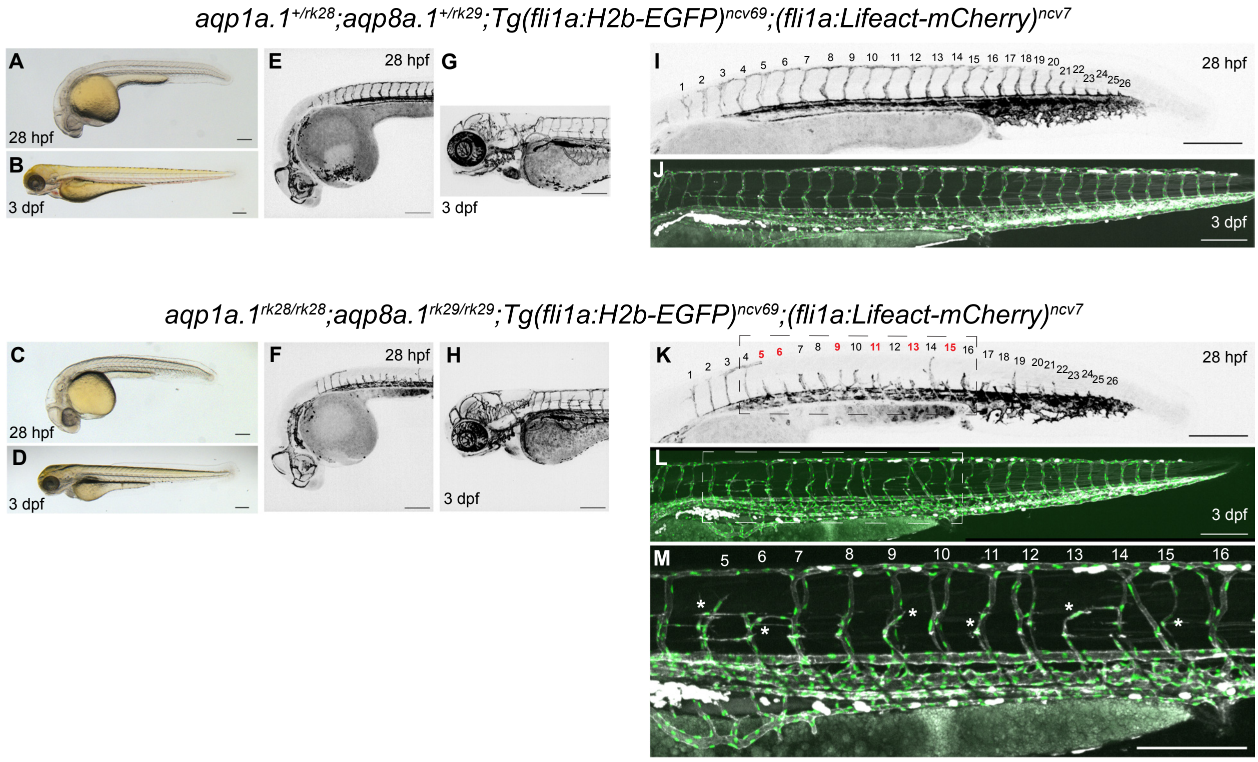

**Figure S8. Phenotype of $aqp1a.1^{rk28/rk28}$;$aqp8a.1^{rk29/rk29}$ double mutant at 28 hpf and 3 dpf.**
The same $aqp1a.1^{+/rk28}$;$aqp8a.1^{+/rk29}$ heterozygote and $aqp1a.1^{rk28/rk28}$;$aqp8a.1^{rk29/rk29}$ homozygote embryos were imaged at 28 hpf and 3 dpf. (**A-D**) Brightfield images of 28 hpf (A, C) and 3 dpf (B, D) embryos showing normal gross morphology of $aqp1a.1^{rk28/rk28}$;$aqp8a.1^{rk29/rk29}$ embryo. (**E-H**) Maximum intensity projections of confocal z-stacks of the brain vasculature of 28 hpf (E, F) and 3 dpf (G, H) embryos. (**I-M**) Maximum intensity projections of confocal z-stacks of the trunk vasculature of 28 hpf (I, K) and 3 dpf (J, L, M) embryos. The trunk and tail were imaged separately and stitched using ImageJ stitching plugin to make an entire image. A magnified image of the boxed areas in K and L is shown in M. In panel K, numbers in red correspond to those sprouts (ISV5, ISV6, ISV9, ISV11, ISV13 and ISV15) that will not form complete ISVs at 3 dpf (asterisks in M). Scale bar, 200 μm.

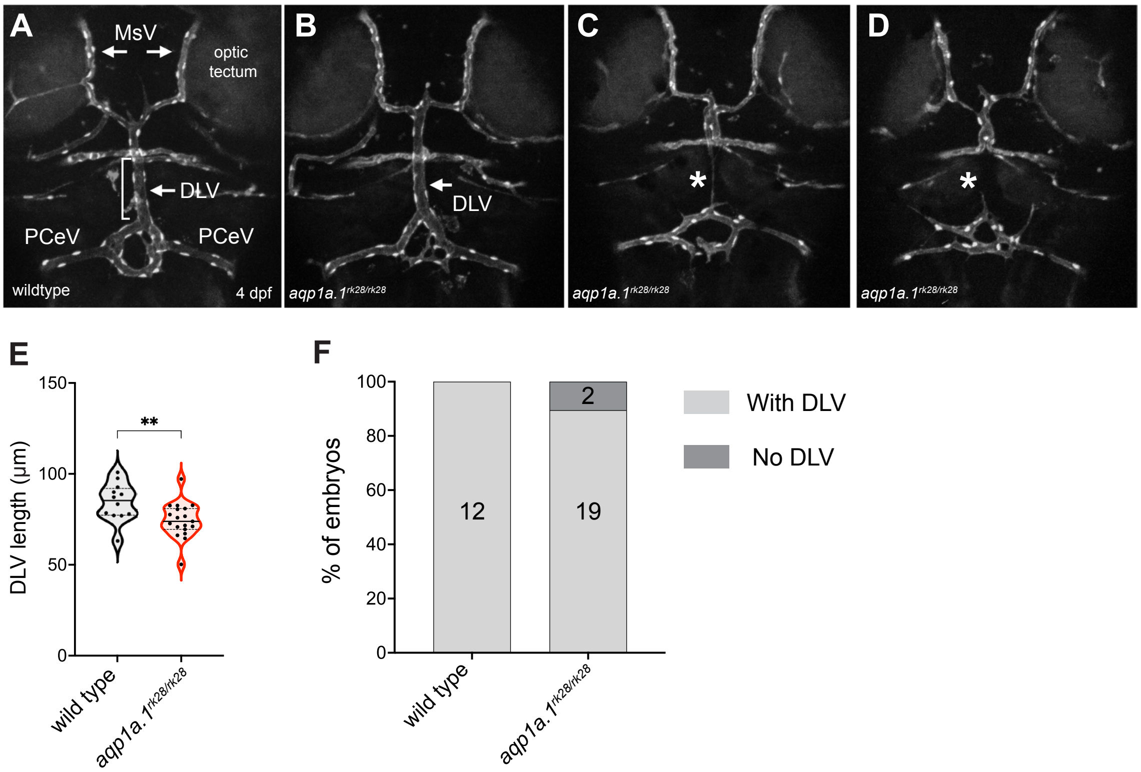

**Figure S9. Cerebral vascular formation defects in *aqp1a.1* mutant zebrafish.**
(**A**) Dorsal view of 4 dpf wildtype cranial vasculature visualized by *Tg(fli1a:H2B-EGFP)*$^{ncv69}$*;(fli1:Lifeact-mCherry)*$^{ncv7}$ expression, showing distinct cranial vessels. (**B-D**) Dorsal view of cranial vasculature of 4 dpf *aqp1a.1*$^{rk28/rk28}$ zebrafish, showing one representative embryo with DLV (B, arrow) and two other embryos lacking the DLV (C and D, asterisks). (**E**) Quantification of DLV lengths of embryos that formed the DLV at 4 dpf ( n=12 wildtype embryos; n=19 *aqp1a.1*$^{rk28/rk28}$ embryos, 2 independent experiments). Statistical significance was determined by unpaired *t*-test; ***p*<0.01. (**F**) Percentage of the embryos of indicated genotype with and without the DLV at 4 dpf (n=12 wildtype embryos, n=21 *aqp1a.1*$^{rk28/rk28}$ embryos, 2 independent experiments). DLV, dorsal longitudinal vein; MsV, mesencephalic vein; PCeV, posterior cerebral vein.

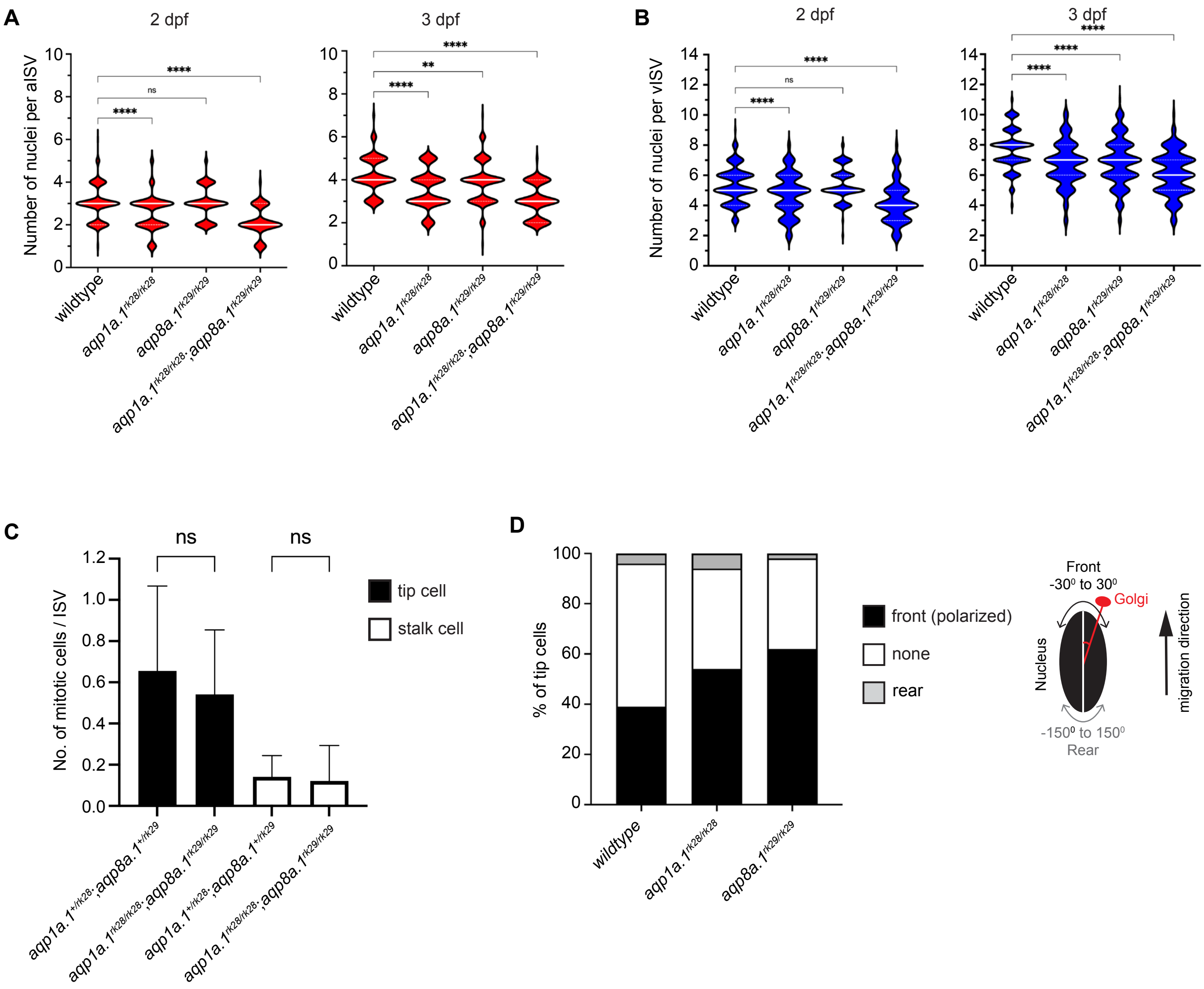

**Figure S10. Reduction in EC number per ISV in *aqp1a.1*$^{rk28/rk28}$*;aqp8a.1*$^{rk29/rk29}$ embryos.**
(**A**) Quantification of EC number in arterial ISVs (aISVs) of wildtype and *aquaporin* mutant embryos at 2 dpf (wild type: n=273 aISVs from 31 embryos; *aqp1a.1*$^{rk28/rk28}$: n=189 aISVs from 21 embryos; *aqp8a.1*$^{rk29/rk29}$: n=84 aISVs from 10 embryos; *aqp1a.1*$^{rk28/rk28}$*;aqp8a.1*$^{rk29/rk29}$: n=65 aISVs from 13 embryos) and 3 dpf (wild type: n=98 aISVs from 10 embryos; *aqp1a.1*$^{rk28/rk28}$: n=86 aISVs from 20 embryos; *aqp8a.1*$^{rk29/rk29}$: n=175 aISVs from 22 embryos; *aqp1a.1*$^{rk28/rk28}$*;aqp8a.1*$^{rk29/rk29}$: n=102 aISVs from 22 embryos). Data collected from 3 (wildtype) and 2 (*aquaporin* mutants) independent experiments. (**B**) Quantification of EC number in venous ISVs (vISVs) of wildtype and *aquaporin* mutant embryos at 2 dpf (wildtype: n=266 vISVs from 31 embryos; *aqp1a.1*$^{rk28/rk28}$: n=143 vISVs from 21 embryos; *aqp8a.1*$^{rk29/rk29}$: n=121 vISVs from 10 embryos; *aqp1a.1*$^{rk28/rk28}$*;aqp8a.1*$^{rk29/rk29}$: n=95 vISVs from 13 embryos) and 3 dpf (wild type: n=121 vISVs from 10 embryos; *aqp1a.1*$^{rk28/rk28}$: n=143 vISVs from 20 embryos; *aqp8a.1*$^{rk29/rk29}$: n=280 vISVs from 22 embryos; *aqp1a.1*$^{rk28/rk28}$*;aqp8a.1*$^{rk29/rk29}$: n=160 vISVs from 22 embryos). Data collected from 3 (wildtype) and 2 (*aquaporin* mutants) independent experiments. (**C**) Quantification of mitotic cells number per ISV in *aqp1a.1*$^{+/rk28}$*;aqp8a.1*$^{+/rk29}$ and *aqp1a.1*$^{rk28/rk28}$*;aqp8a.1*$^{rk29/rk29}$ embryos following live-cell imaging from 21 to 30 hpf (*aqp1a.1*$^{+/rk28}$*;aqp8a.1*$^{+/rk29}$, n=34 ISVs from 5 embryos, 4 independent experiments; *aqp1a.1*$^{rk28/rk28}$*;aqp8a.1*$^{rk29/rk29}$, n=43 ISVs from 9 embryos, 3 independent experiments). (**D**) Quantification of tip cell polarization in wild type (n=79 cells from 10 embryos), *aqp1a.1*$^{rk28/rk28}$ (n=35 cells from 16 embryos) and *aqp8a.1*$^{rk29/rk29}$ (n=90 cells from 9 embryos) embryos at 24-26 hpf. The orientation of the Golgi apparatus relative to the nucleus was assessed to determine cell polarization: the position of the Golgi apparatus within $-30^0$ ~ $+30^0$ angle of the direction of cell migration was considered as polarized. Data collected from 2 independent experiments.
Statistical significance was determined by ordinary one-way ANOVA with Sidak's multiple comparison test (A-B) and unpaired *t*-test (C). ns, $p>0.05$, $^{**}p<0.01$ and $^{****}p<0.0001$. ISV, intersegmental vessel.

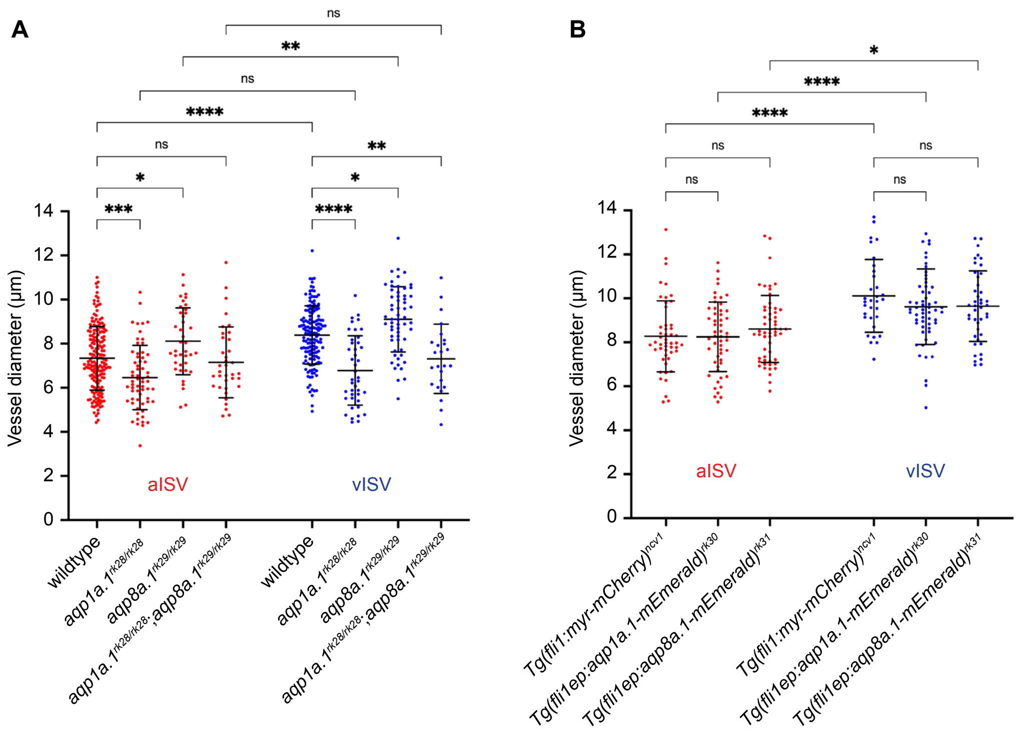

**Figure S11. Depletion of *aqp1a.1* and *aqp8a.1* expression alters vessel diameter.**
(**A**) Quantification of vessel diameter in wildtype and *aquaporin* mutant embryos at 2 dpf (wildtype: n=178 aISVs/148 vISVs from 31 embryos; *aqp1a.1*$^{rk28/rk28}$: n=63 aISVs/42 vISVs from 21 embryos; *aqp8a.1*$^{rk29/rk29}$: n=39 aISVs/59 vISVs from 10 embryos; *aqp1a.1*$^{rk28/rk28}$*;aqp8a.1*$^{rk29/rk29}$: n=36 aISVs/25 vISVs from 13 embryos). Data collected from 3 (wildtype) and 2 (*aquaporin* mutants) independent experiments. (**B**) Quantification of vessel diameter in *Tg(fli1:myr-mCherry)*$^{ncv1}$ (n=49 aISVs/32 vISVs from 8 embryos), *Tg(fli1ep:aqp1a.1-mEmerald)*$^{rk30}$ (n=53 aISVs/58 vISVs from 15 embryos) and *Tg(fli1ep:aqp8a.1-mEmerald)*$^{rk31}$ (n=56 aISVs/43 vISVs from 12 embryos) transgenic embryos at 2 dpf. Data collected from 2 independent experiments. Statistical significance was determined by ordinary one-way ANOVA with Sidak's multiple comparison test. ns, $p>0.05$, $*p<0.05$, $**p<0.01$, $***p<0.001$ and $****p<0.0001$. aISV, arterial intersegmental vessel; vISV, venous intersegmental vessel.

**Supplementary Table 1. Plasmids used in this study.**

| Plasmid name | Plasmid backborne | Method of generation | Reference | Experiment | Comments |
|---|---|---|---|---|---|
| *pMiniT-aqp1a.1* | *pMiniT 2.0 (NEB)* | PCR cloning | This paper | Fig.S1, S4D | Used for riboprobe synthesis and In-fusion cloning |
| *pMiniT-aqp8a.1* | *pMiniT 2.0 (NEB)* | PCR cloning | This paper | Fig.S1, S5D | Used for riboprobe synthesis and In-fusion cloning |
| *fli1ep:aqp1a.1-mEmerald* | *pDestTol2CG2-crybb1:mKate2* | In-Fusion cloning | This paper | Fig.2A | Used to generate *Tg(fli1ep:aqp1a.1-mEmerald)* transgenic line |
| *fli1ep:aqp8a.1-mEmerald* | *pDestTol2CG2-crybb1:mKate2* | In-Fusion cloning | This paper | Fig.2B | Used to generate *Tg(fli1ep:aqp8a.1-mEmerald)* transgenic line |
| *kdrl:mEmerald* | *pMDS6* | In-Fusion cloning | This paper | Fig.5 | *In vivo* cell volume analysis |
| *6xUAS:aqp1a.1-P2A-EGFP-T2A-mKate2-CAAX* | *pMDS6* | In-Fusion cloning | This paper | Fig.6 | Overexpression of Aqp1a.1 protein |
| *fli1ep:nlsEGFP-P2A-mKate2-GM130* | *pMDS6* | In-Fusion cloning | This paper | Fig. S10D | Assessment of tip cell polarization |
| *pDestTolpA2-kdrl:mito-roGFP2-Orp1* | | | Gift from Massimo M. Santoro (University of Turin, Italy) | | Used as a source of *kdrl* promoter |
| *pAc-SP6* | | | Gift from Sergei Parinov (TLL, Singapore) | | Used for *Ac* transposase mRNA synthesis |
| *pCS-TP* | | | Gift from Koichi Kawakami (National Institute of Genetics, Japan) | | Used for *Tol2* transposase mRNA syntesis |
| *pMDS6* | | | Gift from Sergei Parinov (TLL, Singapore) | | Used as a *Ds* transposon source for In-Fusion cloning |
| *pDestTol2CG2-crybb1:mKate2* | | | Gift from Darren Gilmour (EMBL, Heidelberg) | | Used as a backbone for In-Fusion cloning |
| *p5E-fli1ep* | | | Gift from Nathan Lawson (Univ. Massachusetts Medical School) | | Used as a source of *fli1ep* promotor |
| *pCMV:mEmerald-N1* | | | Addgene plasmid #53976 | | Used as a source of mEmerald protein |
| *pDestTol2-fli1ep:mCherry-GM130* | | | Gift from Darren Gilmour (EMBL, Heidelberg) | | Used as a source of GM130 protein |

**Supplementary Table 2. Oligonucleotides used in this study.**

| | |
|---|---|
| Primers for genotyping | |
| aqp1a.1-F1 | CGCCTCCAGATTCATTAGCAGGA |
| aqp1a.1-i1R | GTAAGTGAACTGCTGCCAGTGA |
| aqp8a.1-fwd | GGATCAATTGAGTTGCATAACAGAC |
| aqp8a.1-i1R | CTGTAATGTAGACTTGTAAAGTGGA |
| Primers for the full-length cDNA cloning | |
| aqp1a.1-F1 | CGCCTCCAGATTCATTAGCAGGA |
| aqp1a.1-R1 | CCTGAGGTACATACTGATTCGCTGA |
| aqp8a.1-F | CCGGGCAGAAATCAAAAAT |
| aqp8a.1-R | GCTTGCAATCCTCTTCAGTTC |
| qPCR primers | |
| aqp1a1-qPCRfwd | GGGACTGAATCAAATCCACACAG |
| aqp1a1-qPCRrev | AACCTGATGGCTGTCAGATGT |
| aqp8a1-qPCRfwd | GGAAATCAGTGGTGGTCACTTCA |
| aqp8a1-qPCRrev | TTGTAGTCACAGCTTTGGCAAG |
| gapdh-Fwd | GCTGGTATTGCTCTCAACGATCA |
| gapdh-Fwd | ATGGGAGAATGGTCGCGTATCA |
| hey1-fwd | AAACGTCGCAGAGGGATCAT |
| hey1-rev | CCTGTTTCTCAAAGGCGCTG |
| hey2-fwd | ATTGATGTGGGCAGCGAGAA |
| hey2-rev | TCAATGATCCCTCTCCGCTT |
| dll4-fwd | TGGCCAGTTATCCTGTCTCC |
| dll4-rev | CTCACTGCATCCCTCCAGAC |
| tm4sf18-fwd | CTGGATACTGCTTCCTGATCTC |
| tm4sf18-rev | CAAACAGATACCGTCCCTCAT |
| hAQP1-fwd | CTGGGCATCGAGATCATCGG |
| hAQP1-rev | ATCCCACAGCCAGTGTAGTCA |
| hGAPDH-fwd | GCCACATCGCTCAGACACCAT |
| hGAPDH-rev | TGAAGGGGTCATTGATGGCAACA |
| Oligos for aqp1a.1 and aqp8a.1 sgRNA construction | |
| tracrRNA | AAAAGCACCGACTCGGTGCCACTTTTTCAAGTTGATAACGGACTAGCCTTATTTTAACTTGCTATTTCTAGCTCTAAAAC |
| sgRNAaqp1a.1 | ATTTAGGTGACACTATAGACAGCTGGCCAGCAGACCCGTTTTAGAGCTAGAAATAGCAAG |
| sgRNAaqp8a.1 | ATTTAGGTGACACTATAGATGTCTCCCCCATCGCCCGGTTTTAGAGCTAGAAATAGCAAG |

### Movie legends

**Movie S1. Aqp1a.1 protein enrichment at the leading edge of migrating tip cells.** Live imaging of *Tg(fli1ep:aqp1a.1-mEmerald)*$^{rk30}$ transgenic embryo from 25 to 30 hpf (5 min time interval). 00:00, hour:minutes post fertilization.

**Movie S2. Aqp8a.1 protein enrichment at the leading edge of migrating tip cells.** Live imaging of *Tg(fli1ep:aqp8a.1-mEmerald)*$^{rk31}$ transgenic embryo from 25 to 28 hpf (5 min time interval). 00:00, hour:minutes post fertilization.

**Movie S3. Sprouting angiogenesis in *aqp1a.1*$^{+/rk28}$*;aqp8a.1*$^{+/rk29}$ heterozygote embryos.**

Live imaging of *aqp1a.1*$^{+/rk28}$*;aqp8a.1*$^{+/rk29}$ heterozygote embryos in *Tg(fli1a:H2B-EGFP)*$^{ncv69}$*;(fli1:Lifeact-mCherry)*$^{ncv7}$ double transgenic background from 22 to 30 hpf (7 min time interval). 00:00, hour:minutes post fertilization.

**Movie S4. Defective sprouting angiogenesis in *aqp1a.1*$^{rk28//rk28}$*;aqp8a.1*$^{+rk29/rk29}$ embryos.**

Live imaging of *aqp1a.1*$^{rk28/rk28}$*;aqp8a.1*$^{rk29/rk29}$ mutant embryos in *Tg(fli1a:H2B-EGFP)*$^{ncv69}$*;(fli1:Lifeact-mCherry)*$^{ncv7}$ double transgenic background from 28 to 40 hpf (6 min time interval). 00:00, hour:minutes post fertilization. Arrows point the retraction of tip cell protrusions and the failure of tip cells to invade into the zebrafish trunk from the dorsal aorta.

**Movie S5. Tip cells form an elongated protrusion in the direction of migration during sprouting angiogenesis.**

Live imaging of *aqp1a.1*$^{+/rk28}$*;aqp8a.1*$^{+/rk29}$ heterozygote embryos in *Tg(fli1:Lifeact-mCherry)*$^{ncv7}$ transgenic background from 25 to 30 hpf (7 min time interval). 00:00, hour:minutes post fertilization.

**Movie S6. Defective tip cell leading edge expansion in *aqp1a.1*$^{rk28/rk28}$*;aqp8a.1*$^{rk29/rk29}$ embryos.**

Live imaging of *aqp1a.1*$^{rk28/rk28}$*;aqp8a.1*$^{rk29/rk29}$ mutant embryos in *Tg(fli1:Lifeact-mCherry)*$^{ncv7}$ transgenic background from 25 to 30 hpf (7 min time interval). 00:00, hour:minutes post fertilization.

**Movie S7. The chloride channel SWELL1 promotes EC migration and sprouting angiogenesis.**

Live imaging of *Tg(fli1ep:EGFP-PLC1δPH)$^{rk26}$;(fli1:Lifeact-mCherry)$^{ncv7}$* embryos from 25 to 30 hpf (5 min time interval). Embryos were treated either with 0.05% EtOH or 5 uM DCPIB from 20 to 25 hpf and then imaged. 00:00, hours:minutes post fertilization.

**Movie S8. Additive function of actin polymerization and hydrostatic pressure in driving EC migration and sprouting angiogenesis.**

Live imaging of wild type and *aqp1a.1$^{rk28/rk28}$;aqp8a.1$^{rk29/rk29}$* mutant embryos in *Tg(fli1ep:EGFP-PLC1δPH)$^{rk26}$;(fli1:Lifeact-mCherry)$^{ncv7}$* double transgenic background. Embryos were treated either with DMSO or Latrunculin B from 20 to 28 hpf.
